# CRYPTOCHROME suppresses the circadian proteome and promotes protein homeostasis

**DOI:** 10.1101/2020.05.16.099556

**Authors:** David C.S. Wong, Estere Seinkmane, Alessandra Stangherlin, Aiwei Zeng, Nina M. Rzechorzek, Andrew D. Beale, Jason Day, Martin Reed, Sew Peak Chew, Christine T. Styles, Rachel S. Edgar, Marrit Putker, John S. O’Neill

## Abstract

The daily organisation of most mammalian cellular functions is attributed to circadian regulation of clock-controlled protein expression, driven by daily cycles of CRYPTOCHROME-dependent transcriptional feedback repression. To test this, we compared the circadian proteome and phosphoproteome of wild type and CRY-deficient fibroblast cells. Strikingly, CRY-deficient cells showed a two-fold increase in circadian-regulated proteins, phosphopeptides, and K^+^ transport. This was accompanied by extensive remodelling of the cellular proteome overall, including reduced phosphatase and proteasome subunit expression. These adaptations rendered CRY-deficient cells more sensitive to stress, which may account for their reduced circadian robustness and contribute to the wide-ranging phenotypes of CRY-deficient mice. We suggest that CRY ultimately functions to suppress, rather than generate, daily rhythms in cellular protein abundance, thereby maintaining protein and osmotic homeostasis.

## Introduction

From transcriptional activation and RNA processing, to protein synthesis, folding and degradation, multiple mechanisms operate at every stage of gene expression to ensure that each cellular protein is maintained in a concentration range appropriate to its biological function ^1–3^. Proteome homeostasis is essential for cell viability and osmotic homeostasis, with the correct temporal regulation of protein activity being critical to every biological process—too much or too little at the wrong time underlies most pathological states ^4,5^.

In mammals, cellular physiology is temporally orchestrated around daily cycles that regulate most biological functions and much of the proteome to a circadian rhythm, whereas circadian dysregulation is strongly associated with pathological states ^6^. On any given day, the circadian cycle is expressed by more cells of the human body than the cell division cycle, with daily rhythms in clock-controlled protein abundance thought to be the fundamental basis by which cell biology anticipates and accommodates the predictable demands of day and night ^6,7^. Individual cellular rhythms are synchronised by endocrine cues such as insulin and glucocorticoid signalling, which function *in vivo* to align internal cellular timing with environmental cycles ^8^. Daily rhythms in proteome composition are readily observed *in vivo* and persist in cultured cells under constant conditions *ex vivo* ^9,10^. The average daily variation of rhythmically abundant proteins across a range of cellular contexts is ∼10-20% ^9–12^. There is little direct evidence that such modest variation would necessarily elicit rhythms in protein function, however, given that protein abundance is rarely rate-limiting for protein activity under physiological conditions ^13–16^.

Circadian regulation of a protein’s abundance is most frequently attributed to cycling transcription of the encoding gene, despite recent investigations having revealed that mRNA and protein abundances correlate quite poorly, and that post-transcriptional and post-translational regulatory processes are at least as important ^17–21^. Global transcriptional oscillations are facilitated by daily cycles of transcriptional repression and derepression effected by products of the *Period1/2* and *Cryptochrome1/2* genes, and fine-tuned by various auxiliary but non-essential transcriptional feedback mechanisms ^17,18^. The transcription of *Period*, *Cryptochrome* and other genes is stimulated by complexes containing the activating transcription factor BMAL1. The stability, interactions, and nucleocytoplasmic shuttling of the encoded PER and CRY proteins is regulated post-translationally until, many hours later, they repress the activity of BMAL1-containing complexes in a transcriptional-translation feedback loop (TTFL).

Within the TTFL circuit, CRY proteins are the essential repressors of BMAL1 complex activity ^22,23^, whereas, PER proteins play critical signalling and scaffolding roles, required for the nuclear import and targeting of CRY to BMAL1-containing complexes ^23^. CRY proteins also function as adaptors for the recruitment of E3 ubiquitin ligase complexes that target many proteins, including transcription factors, for ubiquitin-mediated proteolysis ^24^. According to the current widely-accepted paradigm therefore, CRY-mediated transcriptional feedback repression is indispensable for the cell-autonomous circadian regulation of gene expression ^25–29^, with resultant rhythms in clock-controlled protein abundance driving the daily co-ordination of cellular activity ^30^.

Critically though, CRY-deficient cells and tissues remain competent to sustain circadian timing in the absence of TTFL function ^31–33^, with a mechanism that is dependent on casein kinase 1 and protein degradation, as in wild type controls, as well as in naturally anucleate red blood cells ^34–36^. Similarly, circadian oscillations persist in cells and tissue slices lacking BMAL1 ^37,38^. Since BMAL1 and CRY proteins are required for normal circadian regulation *in vivo*, and also required for TTFL function, it was thought that the complex phenotype of BMAL1- or CRY-deficient mice arises because circadian timekeeping is crucial for cellular and organismal physiology more generally ^39,40^. The emerging observation that BMAL1 and CRY-deficient cells and tissues are competent to sustain certain elements of circadian regulation challenges this interpretation and suggests an alternative hypothesis: that some or all phenotypic complexity associated with deletion of a ‘clock gene’ arises from additional functions of the encoded protein beyond the canonical TTFL model. Supporting this, several recent reports imply that various pathophysiological consequences of CRY-deficiency might be directly attributable to the absence of CRY function rather than impairment of circadian transcriptional regulation ^12,41–44^. Overall then, current evidence suggests that whilst CRY-mediated circadian transcriptional feedback repression is crucial for cycling TTFL activity, rhythmic robustness and co-ordinating outputs ^7,23,26,28,29,36^, it is not essential for the cell-intrinsic capacity to maintain daily timekeeping ^33^.

To test this alternative hypothesis, and thereby gain insight into how CRY confers rhythmic robustness, we investigated the molecular consequences of CRY deficiency on cellular protein expression over several days using an unbiased whole cell (phospho)proteomic strategy. Our findings reveal a crucial role for CRY in the maintenance of protein and osmotic homeostasis, with CRY-deficient cells exhibiting a chronic stress-like state that may underlie their reduced robustness as well as the complex and numerous consequences of CRY deletion in mice and their tissues.

## Results

### Cell-autonomous rhythms in the proteome and phosphoproteome persist in the absence of CRY

To understand the proteomic and phosphoproteomic consequences of CRY deletion, we used confluent primary mouse fibroblasts, a classic model of cellular circadian timekeeping where contact inhibition prevents any interference from the cell division cycle ^9^. Wild-type (WT) and CRY1^-/-^; CRY2^-/-^ (CKO) mouse fibroblasts in cell culture were isolated from otherwise isogenic mice and synchronised using daily temperature cycles then sampled under constant conditions (Figure 1A). Quantitative proteomics detected over 6000 proteins and around 4000 phosphopeptides in both cell types. We were surprised to find that, averaging across the time course, the overall abundance and relative phosphorylation of most detected proteins was significantly altered in CKO cells compared with WT controls (Figure 1B, C), by an average of 21%. This striking CRY-dependent change in overall proteome composition was similar to previous findings in CKO mouse liver ^12^ (Figure S1A).

**Figure 1:**
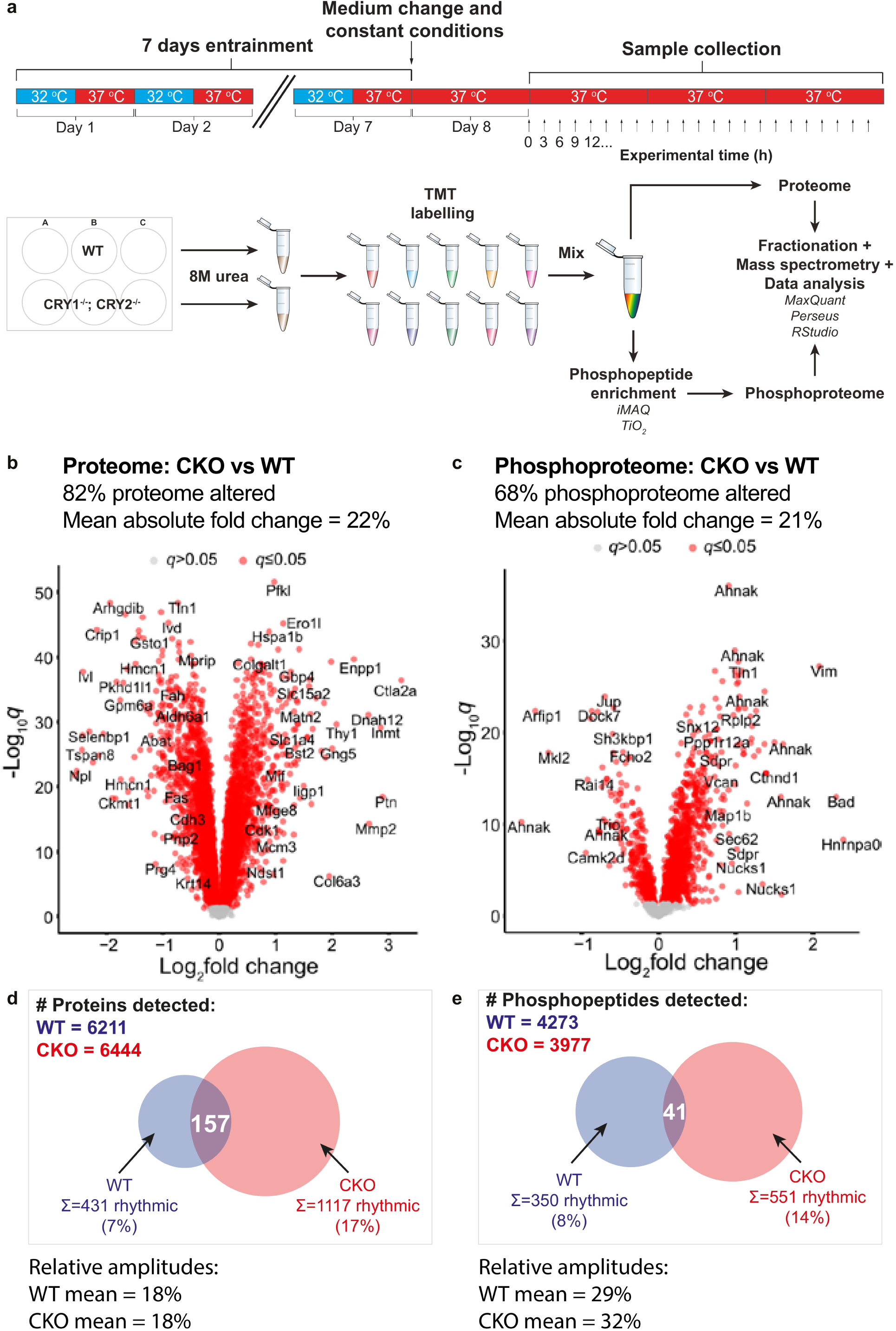
Cell-autonomous rhythms in the proteome and phosphoproteome persist in the absence of CRY. **a)** An overview of the proteomics experimental workflow. Samples were taken every 3 hours for 3 days in constant conditions, starting 24 h after medium change (“Experimental time 0 h”). **b), c)** Volcano plots showing the fold change in average expression of all proteins and phosphopeptides respectively, in CKO cells compared to WT (q = Benjamini-Hochberg corrected p-value, n= 24 time points over 3 days). Statistically significant changes (q ≤ 0.05) are shown in red. Some proteins are labelled as space allows. **d), e)** Venn diagrams showing the numbers of rhythmic proteins and phosphopeptides respectively, in WT cells and CKO cells, with the overlaps annotated. Mean relative amplitude of rhythmic (phospho)peptides are also provided.

In our cellular time course, as expected ^26,28,45^, CRY1 was selectively detected in WT, but not CKO cells, and displayed a ∼24h rhythmic abundance profile across the time course with delayed phase relative to a PER2 luciferase reporter (PER2::LUC) recorded from parallel replicate cultures (Figure S1B). Heatmaps visualising the rhythmic proteins are shown in Figure S1D/E; with examples of rhythmic and arrhythmic (phospho)proteins shown in Figure S1F/G. Based on estimates of intrinsic noise of gene expression ^46–50^, we chose a threshold of 10% relative amplitude to define biological significance for protein abundance oscillations; no such studies were available for protein phosphorylation.

In WT cells, 7% of detected proteins and 8% of detected phosphopeptides showed significant circadian abundance rhythms. Unexpectedly, 17% of detected proteins and 14% of phosphopeptides in CKO cells were rhythmically abundant (Figure 1D, E). Using an independent statistical tool to test for rhythmicity (eJTK cycle), we found that similarly, more rhythmic species were present in CKO cells compared with WT, and with small overlap (Figure S1C).

### CRY suppresses the cell-autonomous rhythmic proteome and phosphoproteome

Amongst the minority of proteins that were rhythmically abundant in both genotypes (Figure 1D), there was a modest but significant increase of median relative amplitude in CKO cells compared with WT (Figure 2A). We calculated the mean abundance of each detected protein over time and divided them into deciles of abundance. We then plotted the proportion of proteins within each decile that were rhythmic against the abundance mid-point of each decile. In this way we found that more abundant proteins were more likely to be rhythmic than less abundant proteins, but crucially, this relationship was stronger for CKO cells compared to WT cells (Figure 2B). Although part of this correlation may be due to preferential and more accurate detection of oscillating high abundance proteins, this explanation cannot account for the different relationships in WT and CKO cells. As with proteins, we found that CKO rhythmic phosphopeptides were significantly increased in relative amplitude and abundance compared with WT (Figure 2C, D). Therefore, the TTFL-independent circadian rhythm seen in CKO cells drives oscillations of higher relative amplitude and acts preferentially towards more abundant proteins and phosphopeptides.

**Figure 2:**
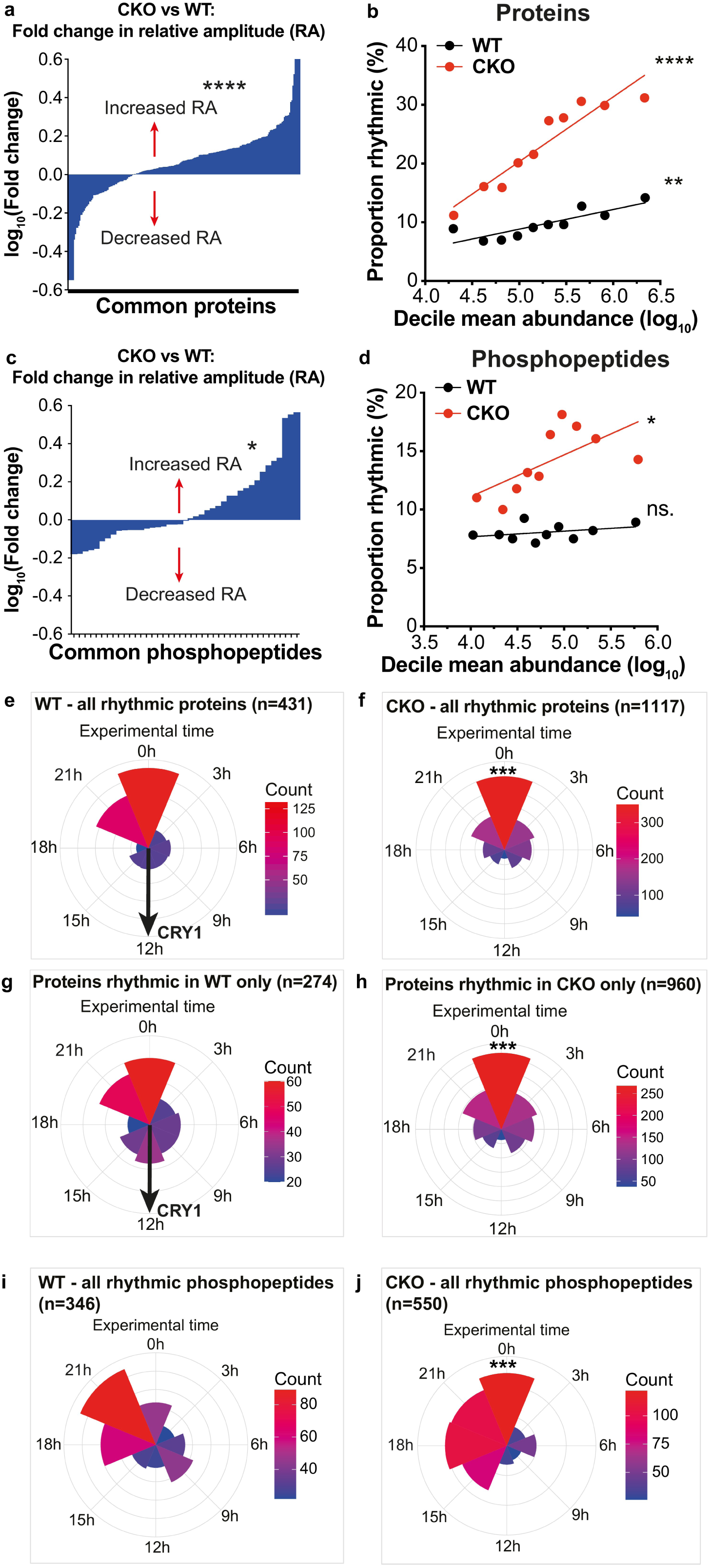
CRY regulates relative amplitude and phase of the rhythmic proteome and phosphoproteome. **a)** Fold-change in relative amplitude (RA) was calculated for each of the proteins found to be rhythmic in both genotypes by RAIN analysis (i.e. no RA cut-off). The mean fold change (log) in RA was an increase of 14% in CKO cells compared to WT cells (One sample t-test, p<0.0001). **b)** For each genotype, all proteins were divided into 10 deciles of equal number, ranked by abundance. The mean abundance of each decile was plotted against the proportion of the decile that was rhythmic. Linear regression lines are shown for each genotype, and the slopes were significantly non-zero (F test, WT p=0.0016, CKO p<0.0001). The slopes were also significantly different to each other (F test, p=0.0001). **c)** Fold-change in relative amplitude (RA) was calculated for each of the phosphopeptides found to be rhythmic in both genotypes by RAIN. On average, the RA was increased in CKO cells compared to WT cells by 6% (One sample t-test, p<0.05). **d)** The same analysis in b) was carried out for phosphopeptides. The slopes were significantly non-zero in CKO but not WT (F test, WT p=0.3, CKO p=0.04). The slopes were also significantly different to each other (F test, p<0.0001). **e, f)** Circular histograms showing the number of proteins at each rhythmic phase. Phase is defined and estimated by RAIN, as the time of the first predicted peak in a 24-hour period. Concentric circles represent the counts scale, with the outermost circle marking the upper end of the counts. The distributions of rhythmic proteins in CKO cells were significantly different to WT cells (left, p<0.001, Watson’s two-sample test). **g), h)** This was also the case for the proteins rhythmic in only one genotype (right, p<0.001, Watson’s two-sample test). **i), j)** Circular histograms showing the number of phosphopeptides at each rhythmic phase. The distributions of rhythmic phosphopeptides in CKO cells was significantly different to WT cells (p<0.001, Watson’s two-sample test).

The phase distribution of rhythmic proteins in both genotypes was clustered at experimental time 0 h (Figure 2E, F), when PER2::LUC activity in wild type cells was maximal (Figure S1A). This phase clustering was also true for proteins that were only rhythmic in CKO cells (Figure 2H), and for most proteins that were only rhythmic in WT cells (Figure 2G). A subset of proteins and phosphopeptides that were rhythmic in WT only, including CRY1, were clustered at a later circadian phase (Figure 2G). This cluster was not present in proteins and phosphopeptides that were rhythmic in CKO only (Figure 2H). In WT cells we observed clustering of protein phosphorylation to an earlier phase (by 3 h) compared to rhythmically abundant proteins and PER2::LUC (Figure 2I) with higher relative amplitude (Figure 1E); this phase relationship has strong similarities with observations in mouse liver *in vivo* ^51^. Whilst the same was true for CKO cells, there was a much broader distribution of phase among rhythmic phosphopeptides across 9 h that preceded peak rhythmic protein abundance (Figure 2J).

These observations demonstrate that cell-autonomous rhythmic regulation of protein abundance and phosphorylation occurs independently of CRY. Instead, the primary role of CRY appears to be two-fold. First, consistent with its function as a rhythmic repressor of gene expression, CRY suppresses the amplitude of cellular protein and phosphopeptide abundance rhythms. Greater amplitudes were observed in the absence of CRY, coupled with more than twice the number of rhythmic proteins detected in WT cells, and with most rhythmic CKO (phospho)proteins not being detectably rhythmic in WT cells (Figure 1D, E). Secondly, in WT cells, CRY regulates the phase of rhythmicity for a sub-population of proteins and phosphopeptides whose abundance peaks around the same phase as CRY1 and in antiphase with the majority of the rhythmic (phospho)proteome.

Overall, we observed that a remarkable 82% of detected proteins in CKO cells were altered in abundance compared with WT (Figure S2A, Abundance up + down), suggesting that the respective synthesis and degradation rates of many proteins are altered by CRY deficiency. We postulated that CRY deletion might unmask rhythmicity in proteins that normally exhibit matched rhythms of synthesis and degradation in WT cells, i.e. constitutively present with rhythmic turnover. Supporting this, there was a significant association between changes in average protein abundance and changes in rhythmicity: we found that proteins that changed in overall abundance were more likely to change in rhythmicity: from arrhythmic to become rhythmic, or *vice versa* (Figure S2A). In this case, our observations would indicate that the cell-autonomous circadian clock regulates ∼ 30% of the fibroblast proteome (Figure S2A, green + blue + brown), with CRY primarily acting to suppress rhythms in the abundance of most of these proteins (Figure S2A, green).

In fact, we found that there was greater overall variation in abundance of proteins detected in CKO cells (Figure S2B), suggesting that CRY may suppress changes in protein levels more generally. To verify this finding using an independent dataset, we calculated the fold-change in abundance for proteins detected by Mauvoisin *et al*. from WT and CKO mouse liver samples ^12^. There was greater variation in protein abundance in CKO compared with WT (Figure S2C). Therefore, CRY also functions to suppress variation in protein abundance in mouse liver *in vivo* under diurnal conditions.

To test the effects of CRY deletion on the phosphoproteome, we performed the same analysis with detected phosphopeptides. As with the proteome, we found a strong link between changes in abundance and rhythmicity of phosphopeptides between the two genotypes (Figure S2D), as well as greater variation in phosphopeptide abundance in CKO compared with WT cells (Figure S2E).

We also observed an increase in overall protein phosphorylation in CKO cells (Figure 1C, S3A). We therefore considered whether CRY deletion might impact upon the abundance or activity of protein kinases and phosphatases, which act in dynamic equilibrium to determine the phosphorylation level of each peptide we detected. In our proteomics dataset there was no significant overall change in abundance of protein kinases (Figure S3B). To validate this, we employed the PHOSIDA database ^52,53^ to infer rhythmic kinase activity; and found that targets of kinases typically associated with circadian regulation were not over-represented in WT or CKO rhythmic datasets compared to background, nor in the set of phosphopeptides that were rhythmic in both genotypes (Figure S3C-E). CKO cells did show an overall decrease in abundance of protein phosphatases however (Figure S3B), which may plausibly account for both increased overall phosphorylation and increased rhythmic phosphorylation in CKO compared with WT cells. This would suggest that CRY normally functions to globally supress steady-state protein phosphorylation, as well as rhythms in protein phosphorylation, in part through regulation of phosphatase abundance.

Bringing these observations together we suggest that the synthesis and degradation of many proteins, as well as the phosphorylation and dephosphorylation of many phosphosites, may be coupled through a mechanism that requires CRY to be present. CRY deletion therefore elicits a significant change in dynamic steady-state abundance for most of the (phospho)proteome, including a sub-set of proteins that are normally circadian-regulated, thereby revealing more than twice the number of rhythmic proteins and phosphopeptides detected in WT cells. The altered protein stoichiometries of CKO compared with WT cells and tissues would be described as a state of proteome imbalance if it occurred in the context of cancer cells ^54^ and is normally associated with altered protein homeostasis. That such proteome imbalance is observed in primary cells and tissues suggests the presence of widespread changes in the cellular determinants of proteome composition ^4,5^.

### CRYPTOCHROME regulates proteasome activity and translation rate

We explored potential mechanisms for the widespread proteome changes in CKO cells by searching our dataset for key factors regulating protein synthesis and degradation. We did not interrogate the transcriptome of CKO cells, which has been adequately addressed elsewhere, and since there is now overwhelming consensus that mRNA abundance is poorly predictive of protein abundance ^10–12,55–64^. We found a striking reduction in the abundance of catalytic proteasomal subunits (Figure 3A) which we validated by western blot and enzymatic assays of proteasome activity (Figure 3B-D, S4A).

**Figure 3.**
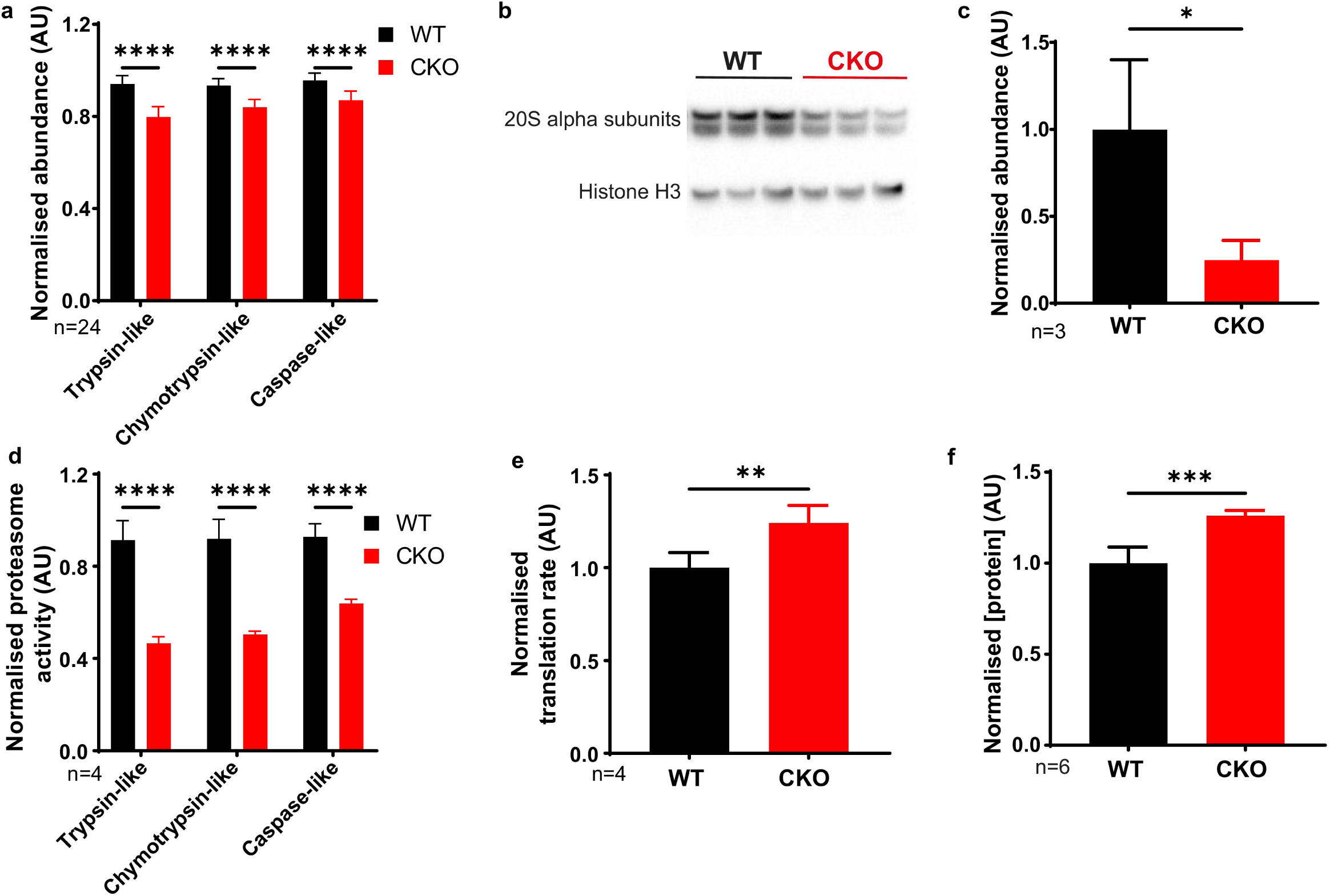
CRYPTOCHROME regulates proteasome activity and translation rate. **a)** From the quantitative proteomics experiment, average abundance of catalytic proteasome subunits was calculated and normalised to WT means. Trypsin-like (β2), chymotrypsin-like (β3) and caspase-like (β1) catalytic subunits are shown. The average was calculated from all 24 time points of the proteomics experiment. Mean±SD, 2-way ANOVA with Holm-Sidak’s multiple comparisons. **b)** Representative Western blot using an antibody that recognises all 7 α subunits of the 20S proteasome, with anti-histone H3 as loading control. **c)** Quantification of the blots in b), using all replicates, normalised to WT mean. Mean±SD, one-tailed Student’s t test with Welch correction – prior prediction from experiment in A) that CKO abundance would be lower. **d)** Proteasome activity measured using the ProteasomeGlo Assay (Promega), normalised to WT means. Mean±SD, 2-way ANOVA with Holm-Sidak’s multiple comparisons. N=6 experiments, representative experiment shown (n=4 technical replicates). **e)** Translation rate was measured using ^35^S-methionine labelling and imaging with phosphor screens. The quantification values were normalised to the total protein concentration as measured using Coomassie stain, and then normalised to WT mean. Mean±SD, Student’s t test with Welch correction. N=3. **f)** Total protein mass per cell in confluent WT and CKO cultures. Cells were grown in two 12-well plates; one was used for cell counting and the other was used for lysis in RIPA buffer prior to protein quantification by BCA assay. Quantification shown, normalised to WT mean. Mean±SD, Student’s t test with Welch correction.

Whilst there was no consistent change in ribosomal subunit abundance, we also observed a significant increase in cytosolic protein synthesis rate by ^35^S-methionine incorporation (Figure 3E, S4B, C), as observed previously by puromycin incorporation ^33^. We considered that the combined effect of reduced proteasomal activity and increased translation rate would affect the overall steady state levels of cellular protein. We found this to be the case, with a modest but significant increase in overall protein per CKO cell compared with WT (Figure 3F).

These cultured CKO cells express more protein than WT cells, with increased translation and decreased proteasomal degradation. Altogether, and in light of previous observations ^65–67^, our data suggest that CKO cells likely maintain a different set point for protein homeostasis, as also occurs in many pathological states ^4,5^.

### Rhythmic regulation of ion transport and protein content is CRY-independent

Mammalian and other eukaryotic cells devote significant resources to ensuring osmotic equilibrium over the plasma membrane, in order to maintain cell volume and prevent protein aggregation ^67–70^. Changes in cytosolic macromolecule content are balanced by compensatory ion transport, primarily mediated by K^+^, the major cellular osmolyte ^67,68^. Two independent lines of evidence suggested that CKO cells would exhibit altered osmotic homeostasis.

First, we searched for rhythmically regulated cellular processes in CKO and WT cells using gene ontology (GO) analysis. Of the rhythmically abundant proteins, ranked GO analysis revealed consistent enrichment for processes associated with ion transport, both in wild type and CKO cells when analysed separately or combined (Figure 4A-C). Interrogation of the proteomics dataset then revealed altered expression levels and increased rhythmic amplitudes of many ion transporters in CKO compared with WT cells (Figure S5A, B). This included several members of the SLC12A family of electroneutral transporters (Figure 4F), that contribute to the maintenance of osmotic homeostasis against variations in cytosolic protein concentration in wild type cells over the circadian cycle ^71^. Second, since many proteins are cytoplasmic, we considered that if CRY normally suppresses rhythms in the abundance of many individual proteins that normally peak around the same circadian phase (Figure 2E-H) then the consequence of CRY deletion would be to increase the overall amplitude of daily rhythms in soluble cytosolic protein content.

**Figure 4.**
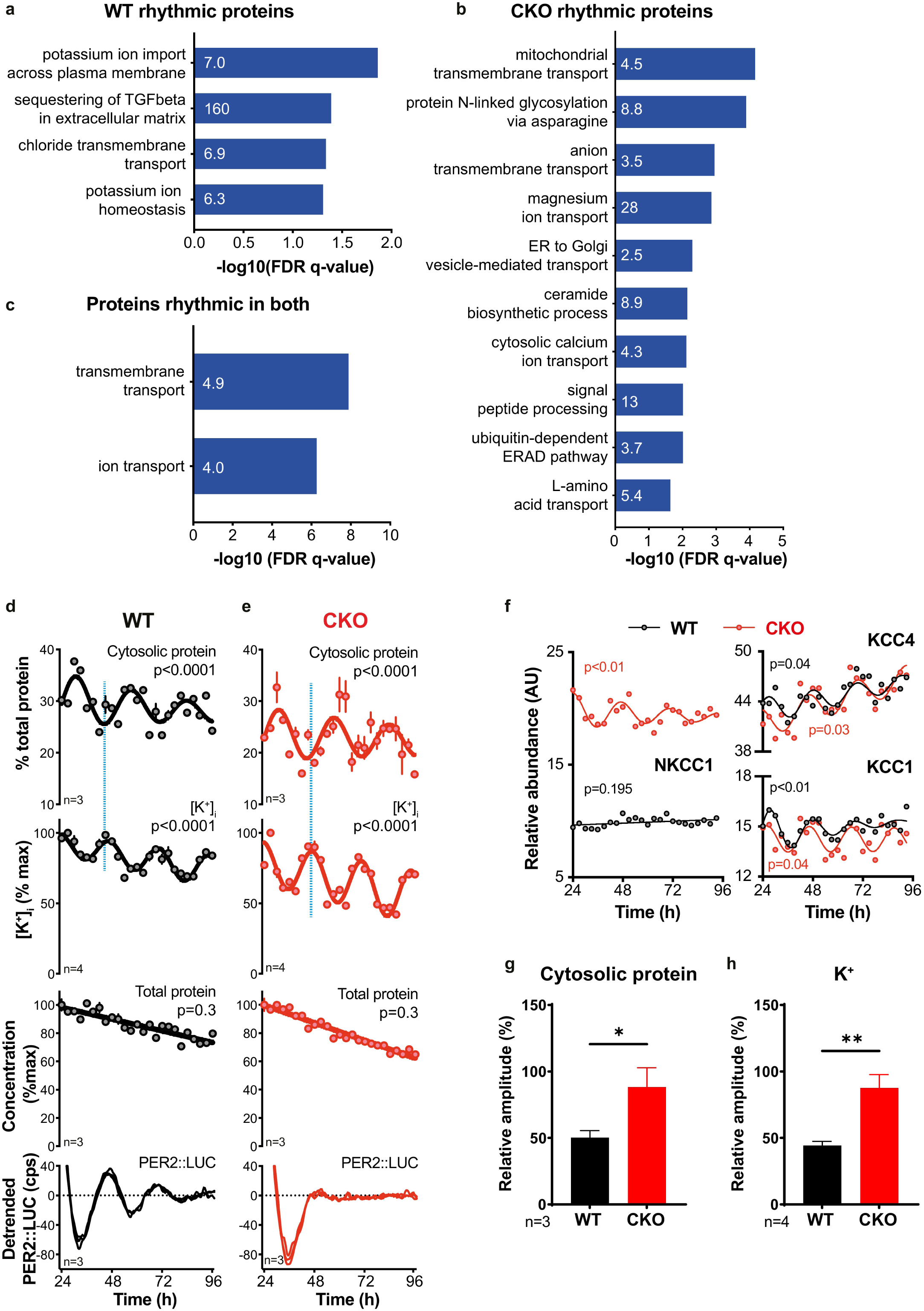
Rhythmic regulation of ion transport in WT and CKO cells. **a)** Gene ontology analysis for rhythmic proteins was carried out using Gene ontology enrichment analysis and visualisation (GOrilla) ^130,131^. Significantly rhythmic WT proteins were compared against background (all proteins identified in the experiment), and the top non-overlapping GO Biological Process terms shown, sorted according to FDR q-value. Fold enrichment is annotated on each bar. The same GO analysis was carried out comparing proteins rhythmic that were rhythmic in CKO cells **(b)** and for proteins that were rhythmic across both genotypes **(c)**. **d), e)** From one time-course experiment, ions, cytosolic proteins and total protein were extracted in parallel samples. The presented experiment is representative of 3 separate time-course experiments that were carried out (N=3). Blue lines highlight the antiphasic relationship between oscillations in cytosolic protein and potassium concentration. Mean±SEM, p-values from RAIN, red lines are fits by a damped cosine compared with a straight line (null hypothesis). Parallel PER2::LUC recordings were also performed and plotted as a phase marker. **f)** Examples of key ion transporters are shown, as detected in the proteomics experiment. P values show the results of an F test comparing fits of damped cosine against straight line. All proteins except WT NKCC1 had RAIN p-values <0.05. **g), h)** Relative amplitudes of cytosolic protein and potassium concentrations oscillations in a) and b) were greater in CKO compared to WT (Student’s t test with Welch correction, mean±SD).

To validate this, we measured the K^+^ content of cells across the circadian cycle. Consistent with previous investigations ^71^, in WT cells, K^+^ and digitonin-extracted cytosolic protein concentrations exhibited antiphasic circadian rhythms (Figure 4D, Figure S5C), with no significant daily variation in total cellular protein. The same was observed in CKO cells (Figure 4E), but with higher relative amplitudes for soluble protein and K^+^ (Figure 4G, H). Considering previous observations ^71^, the higher amplitude cytosolic protein rhythm in CKO cells likely drives the higher amplitude K^+^ rhythms. This is facilitated by increased expression and amplitude of SLC12A transporter activity (Figure 4F), which buffers cellular osmotic potential in response to greater changes in cytosolic macromolecular content over the circadian cycle ^71^.

Previous work has suggested that compensatory K^+^ transport is a fundamental feature of eukaryotic cell biology allowing cells to accommodate changes in cytosolic macromolecular content whilst maintaining osmotic homeostasis ^67,71^. Our present observations suggest that this occurs independently of CRY-mediated transcriptional feedback repression. Indeed, it is plausible that an important role of CRY may be to suppress daily changes in cytosolic macromolecular content and thus maintain osmotic balance more efficiently over the circadian cycle. This would effectively increase the capacity of WT cells to respond appropriately to any acute stimulus that requires change in proteome composition e.g. insulin/IGF-1 signalling ^8^.

### CRY-deficient cells are more sensitive to proteotoxic stress

The viability of CKO cells and mice clearly suggests that they do maintain protein homeostasis overall, despite proteome imbalance and an altered set point for protein homeostasis compared with WT controls. Proteome imbalance typically renders cells more sensitive to stress ^3^. Using ranked GO analysis of overall protein fold-changes compared with WT, we found that the expression of proteins involved in “response to stress” was increased in CKO cells (Figure S6A), suggesting that the proteostasis network in CKO cells is in an activated state ^66,72,73^.

We therefore hypothesised that CKO cells may be more susceptible to proteotoxic stress. To test this, we probed WT and CKO cells for phosphorylation of eIF2α, a well-characterised marker of the integrated stress response (ISR). As a control we treated cells with tunicamycin, which gradually induces the integrated stress response *via* inhibition of secretory pathway protein glycosylation ^74,75^. We observed increased eIF2α phosphorylation in CKO compared with WT cells, at all timepoints (Figure 5A-B). This is strongly indicative of increased proteotoxic stress in these CKO cells.

**Figure 5.**
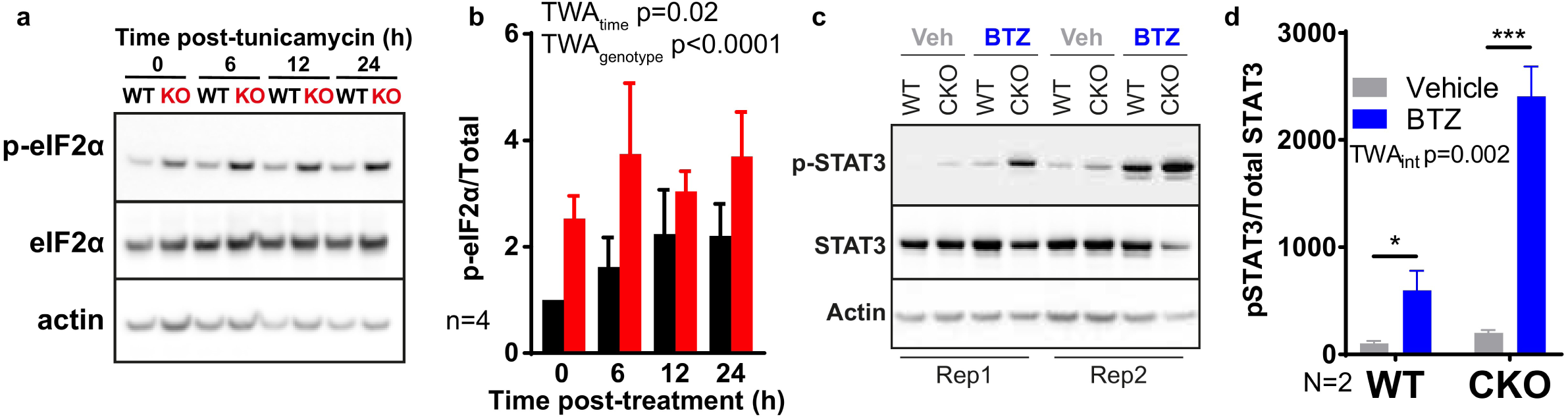
CKO cells and tissues are more sensitive to stress. **a)** WT and CKO cells were treated with 500 nM tunicamycin (TUN) and lysed in RIPA buffer at time points between 0-24 hours afterwards. Western blots were carried out, probing for phosphorylated eIF2α, total eIF2α and actin. Representative blots shown, n=4. **b)** Quantification of all replicates represented in (a), where phosphorylated eIF2α is normalised to total eIF2α. Mean±SD. 2-way ANOVA (TWA) with Holm-Sidak’s multiple comparisons. **c)** Mouse lungs were collected 7 hours after transition to the light phase, 5 hours after intraperitoneal injection of mice with bortezomib (BTZ, 2.5 mg/kg) or vehicle (Veh, 1% DMSO in sterile PBS). Tissues were lysed in RIPA buffer and western blots were performed, probing for phosphorylated STAT3, total STAT3 and actin. All replicates shown, N=2 mice for each condition. **d)** Quantification of the blots shown in (c), where phosphorylated STAT3 is normalised to total STAT3. Mean±SD. 2-way ANOVA (TWA) with Holm-Sidak’s multiple comparisons.

We note that in most cases eIF2α phosphorylation is known to acutely suppress translation ^72^, yet we observed a net increase of protein synthesis in CKO cells (Figure 3C, S4D-E). This may be reconciled by our observation that density regulated protein (DENR) and eIF2A, additional alternative subunits of the translation initiation complex, were significantly upregulated in CKO cells compared with WT (Figure S6B-C); and suggests a mechanism whereby chronically stressed CKO cells may overcome increased basal eIF2α phosphorylation, that normally only occurs under acute stress ^76–79^.

We next sought to test whether increased stress was associated with CRY-deficiency in mouse tissues. STAT3 is a transcription factor that whose phosphorylation on Tyr705 is a well-established inflammatory marker for both chronic and acute cellular stress *in vivo,* across a range of cell types ^80–82^. We therefore probed for phosphorylated STAT3 in mouse lungs 5 hours after intraperitoneal injection of bortezomib (BTZ, proteasome inhibitor) or vehicle (control). Compared with WT tissue, we observed increased STAT3 phosphorylation in control CKO mouse lungs, which was further elevated above WT upon BTZ treatment (Figure 5C-D). This shows CRY-deficiency is associated with increased stress and increased sensitivity to stress in mouse tissue, as well as cultured primary cells.

### Proteotoxic stress impairs rhythmic robustness

Daily rhythms of PER2 activity in CKO cells (as measured by PER2::LUC bioluminescence) are less robust than in WT cells, being more variable in their expression and damping more rapidly ^33^. At the outset of this investigation, we asked how CRY confers increased robustness upon cellular circadian rhythms observed in WT cells, and found evidence suggesting that proteome imbalance in CRY-deficient cells renders them more sensitive to stress. Noting that several different cellular stressors (reductive, oxidative, metabolic, transcriptional inhibition) have previously been reported to reversibly attenuate the expression of cellular circadian rhythms ^83,84^, our observations suggested the hypothesis that the altered set point of protein homeostasis in CKO renders them more susceptible to stress, which in turn leads to their reduced rhythmic robustness. This informed the prediction that circadian rhythms in WT cells would be less robust, and therefore damp more rapidly, when subject to chronic proteotoxic stress. To test this we monitored PER2::LUC rhythms in WT cells where proteotoxic stress was elicited by sustained inhibition of three different pathways: with epoxomicin (proteasome inhibitor), tunicamycin (unfolded protein ER stress response) or radicicol (HSP90 inhibitor). Consistent with our hypothesis, we observed significantly increased damping rate in treated cells compared with controls, which was reversible upon drug removal by a media change (Figure 6A-F). Thus, chronic proteotoxic stress is sufficient to impair robustness of circadian rhythms, and in principle this may contribute to the reduced robustness and timekeeping fidelity of CKO cells and tissues, as well as diverse CKO mouse phenotypes (Figure 6G, S6D-G, S7).

**Figure 6.**
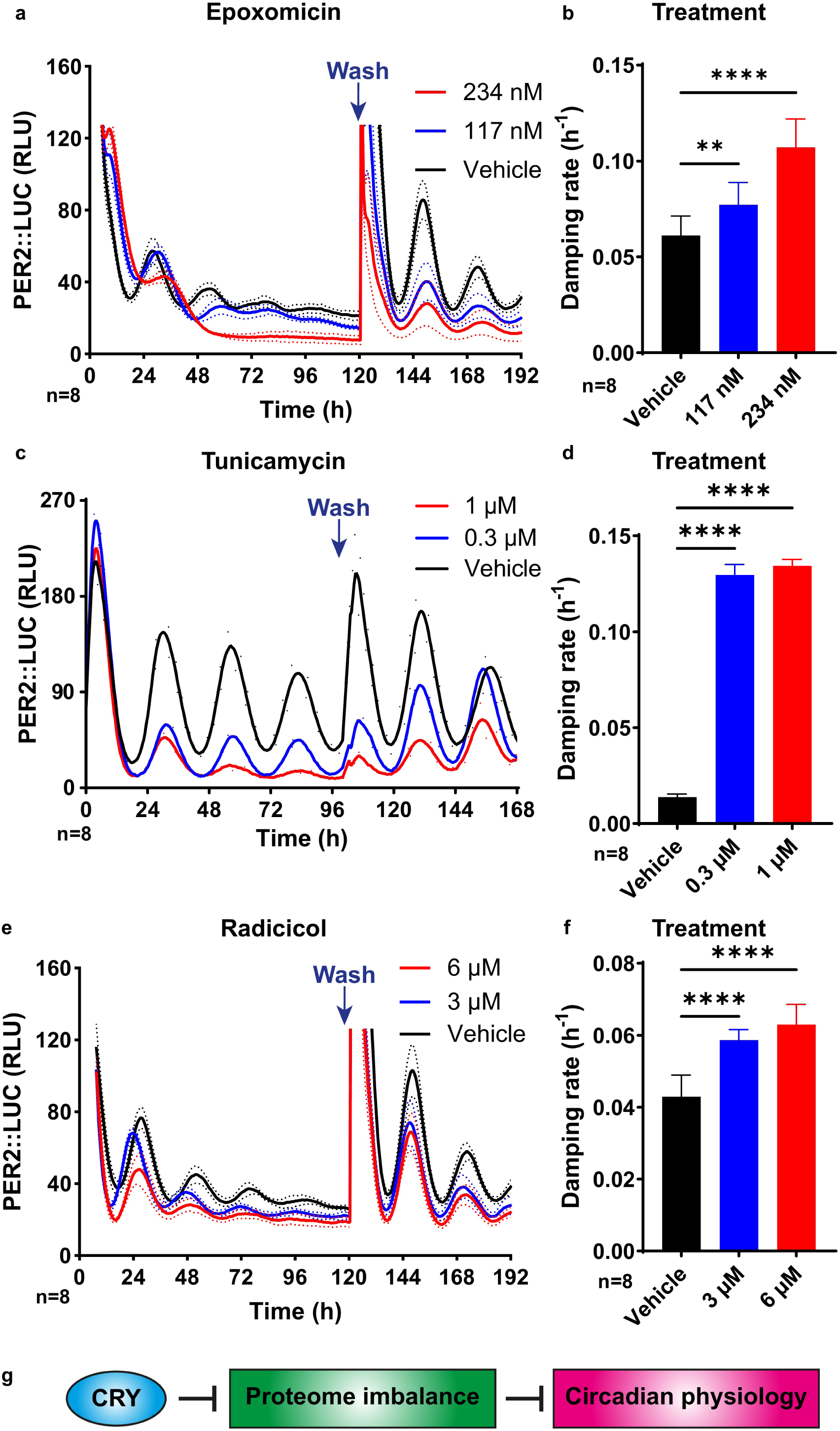
WT cells recapitulate the CKO phenotype upon induced chronic stress. **a)** WT PER2::LUC cells were treated with epoxomicin or vehicle control. Blue arrows show the time points where drug was washed off. Mean±SD. **b)** Quantification of damping rate of the PER2::LUC recordings shown in a), for the duration of drug treatment. Mean±SD. **c)** WT PER2::LUC cells were treated with tunicamycin or vehicle control. Mean±SD. **d)** Quantification of damping rate of the PER2::LUC recordings shown in c), for the duration of drug treatment. Mean±SD. **e)** WT PER2::LUC cells were treated with radicicol or vehicle control. Blue arrows show the time points where drug was washed off. Mean±SD. **f)** Quantification of damping rate of the PER2::LUC recordings shown in e), for the duration of drug treatment. Mean±SD. **g)** Schematic diagram: altogether our data suggests that in the absence of CRY, proteome imbalance disrupts circadian rhythmicity of physiology. See text for details.

## Discussion

### CRY defends cellular homeostasis, which is permissive for robust circadian rhythms

In order to understand the reduced robustness of circadian rhythms in CRY-deficient cells we used several complementary mass spectrometry techniques. We found the abundance of most cellular proteins and protein phosphorylation, as well as the major cellular osmolyte (K^+^), to be profoundly perturbed by CRY-deficiency. This was associated with adaptations such as reduced proteasome activity and phosphatase abundance that alter the relative rates of protein synthesis/degradation, and phosphorylation/dephosphorylation, respectively. This results in altered osmotic homeostasis and a state of proteome imbalance compared with WT cells. Proteome imbalance is established as predisposing cells to stress ^54,85,86^ and is therefore very likely responsible for the increased proteotoxic stress and sensitivity to stress observed in CKO cells. Increased proteotoxic stress by three different mechanisms was sufficient to impair robustness of WT cellular circadian rhythms. Our data therefore suggest that proteome imbalance and associated stress are most likely responsible for the impaired timekeeping fidelity and reduced robustness of CKO cells and tissues *ex vivo*.

CRY proteins are well-characterised as rather promiscuous transcriptional repressors ^42,45,87^ as well as being selective E3 ubiquitin ligase adaptors ^24,88^. Considering the diverse phenotypes, pathologies and proteome changes reported for other E3 ligase adaptor and transcriptional repressor knockout mice ^89–93^, it is not surprising that the deletion of both *Cryptochrome* genes results in a similarly diverse range of phenotypes; especially when considering the very many identified targets and interacting proteins of CRY1 and CRY2 ^24,42,45,87,88^.

Proteome imbalance is already very strongly, and in some cases causally, linked to a range of pathological conditions including chronic inflammation, various cancers, metabolic disorders and neurodegeneration ^4,54, 94–99^. It is therefore quite plausible that proteome imbalance underlies the diverse range of phenotypes exhibited by CKO mice, including impaired body growth ^100,101^, increased susceptibility to multiple cancers ^102–105^, chronic inflammation ^106–108^ and dysregulated insulin secretion/fat deposition on high fat diets ^109^. Indeed, during the course of this research we observed that not only do CKO mice eat more and gain less weight than their isogenic WT counterparts, but they also succumbed to spontaneous mortality/morbidity with a much higher frequency (Figure S6D-G).

### CRY suppresses circadian rhythms in protein abundance and osmotic balance

We also noticed that the profound alteration of cellular (phospho)proteome and ionic composition of CKO cells included a complete reorganisation of the subset of proteins and phosphoproteins that are subject to circadian regulation. To our surprise, roughly twice as many proteins and phosphopeptides were rhythmic in CKO cells compared with WT, but only a small minority of rhythmic species were common to both genotypes. We also found that significant changes in the overall abundance of a given protein or phosphopeptide were associated with a change in rhythmicity. The parsimonious interpretation of our findings is that, directly or indirectly, CRY normally functions to suppress rhythms in the abundance of a substantial proportion (19%) of detected cellular proteins, possibly by coupling rhythms of their synthesis and degradation to maintain a dynamic steady state with rhythmic flux (Figure S2A). Removing CRY unmasks the circadian rhythm for these proteins, with most (>80%) changing in overall abundance, due to the new steady state equilibrium that results from a change in the average rate of protein synthesis relative to degradation.

Importantly, we did observe a smaller proportion of cellular proteins (6% total), upon which CRY normally confers rhythmic abundance and which lose that rhythm when CRY is absent; consistent with the more canonical view of CRY function within the TTFL model for circadian rhythms. The majority of these proteins detected (>80% of 6%) also changed in overall abundance when CRY is absent, achieving a new equilibrium concentration for the same reason as those proteins where rhythmic abundance is suppressed by CRY (Figure S2A).

The hypothesis that CRY proteins primarily function to suppress, not generate, variation in protein abundance is further supported by *post-hoc* analysis of published data from mouse liver collected under diurnal cycles and quantified by an independent method ^12^. If this hypothesis is correct, it informs the explicit prediction that, in wild type cells, a much greater proportion of proteins are subject to phase-coherent circadian regulation of synthesis/degradation than is apparent from their steady state abundance (Figure S7B). The same logic applies for protein (de)phosphorylation, and these predictions will be a major focus of our future investigations.

We also found that, compared to WT, CKO cells have increased protein abundance and reduced K^+^ levels overall, as well as higher amplitude rhythms of cytosolic protein and K^+^. Changes in soluble protein concentration require stoichiometrically larger changes in ion concentration to maintain osmotic homeostasis ^69^. We therefore suggest that CKO cells may have an impaired ability to buffer changes in intracellular osmolarity, particularly in response to external stimuli, which may contribute to their increased sensitivity to stress ^110,111^ and attenuated but prolonged response to growth factor stimulation ^8^. Overall, our findings have significant implications for the role of CRY proteins specifically, and the canonical TTFL more generally, as well as opening several important avenues for future investigation.

### The utility of circadian rhythms for cellular proteostasis

It is frequently suggested that the adaptive advantage of cellular circadian clocks is to anticipate the differential demands of day and night, by turning on genes to accommodate the anticipated increase in demand for the activity of the encoded protein ^112^. Protein synthesis is the most energetically expensive process that most cells undertake however, and multiple mechanisms exist to inactivate and sequester proteins that are not required ^1,86^. Moreover, recent reports favour the view that changes in cellular transcriptomes function to buffer cellular proteomes, not perturb them ^19,20,113^. Rather than synthesise proteins as and when needed, it makes evolutionary sense that cells would expend energy to ensure a constant abundance of most proteins, which could then be mobilised on demand.

However costly though, damaged/misfolded proteins and activated signal transducers do need to be degraded and replaced to avoid deleterious consequences, such as aggregation and sustained pathway activation. We therefore hypothesise that, rather than rhythm generation, a fundamental advantage conferred on mammalian cells by the daily regulation of CRY activity, within and beyond the canonical TTFL circuit, is the temporal consolidation of proteome renewal, matching synthesis and degradation rates to keep protein concentrations constant and maintain protein homeostasis overall. Indeed, synchronised daily increases in protein synthesis and degradation are likely to be a prerequisite for the efficient assembly of macromolecular protein complexes ^114^. Supporting this hypothesis, temporal consolidation of proteome renewal has been observed in yeast cells during their metabolic cycle, a biological oscillation that shares many key features with circadian rhythms in mammalian and other eukaryotic cells ^67^. This hypothesis leads to the direct prediction that macromolecular complex assembly will be less efficient and more energetically expensive in cells and tissues that lack the capacity for daily regulation of gene expression cycles, such as Cry1/2, Per1/2 and Bmal1-knockouts.

### Caveats to our findings

In this investigation, we specifically addressed how the steady state circadian cellular (phospho)proteome adapts to CRY deficiency in order to understand the impairment to cellular timekeeping. For this reason, we did not investigate transcriptional regulation in CKO cells, which we believe has been adequately characterised in excellent previous work ^25,26,28,33, 115–118^. Moreover the generally poor correlation between changes in transcript abundance with protein activity ^19,119–122^ means that causal relationships cannot be reliably inferred in any case. Thus, we do not exclude that circadian regulation of transcription occurs in CRY-deficient cells, indeed we have observed rhythms in the activity of the *Nr1d1* promoter in CKO cells, albeit under very specific culture conditions with <5% the amplitude of WT controls ^33^. Rather, the increased number and amplitude of protein abundances and phosphorylation we observed in CRY-deficient cells simply cannot be attributed to canonical CRY/PER-mediated transcriptional feedback repression, since CRY is not present and PER does not repress BMAL1 complexes without CRY ^22,23^. We also have not addressed the nature of the post-translational mechanism postulated to generate circadian rhythms in mammalian cells, which is discussed elsewhere ^7,33^.

A further caveat to our study is that, for technical reasons, the primary fibroblasts used for (phospho)proteomics came from a single, but otherwise isogenic, WT and CKO mouse. In our previous work, we have demonstrated circadian rhythms in many independently generated CKO fibroblast lines and tissues, isolated from many different mice ^33^. Furthermore, analysis of published mouse liver data provides an independent validation of our finding that CRY primarily functions to supress, not generate, daily protein variation. Moreover, the null hypothesis for this aspect of our analysis was either no rhythms or severely attenuated rhythms in CKO cells, as opposed to higher amplitude rhythms in more proteins, which was subsequently validated by independent methods in separate experiments (cytosolic [protein], & [K^+^]). Therefore, the most parsimonious interpretation of our findings is that CRY functions primarily to oppose daily protein abundance rhythms, rather than generate them, and thereby buffer cellular protein homeostasis over each daily cycle. We are not aware of any evidence to contradict this interpretation and have no reason to believe that there was anything special about the particular primary CKO fibroblasts used in this study.

It is quite plausible, however, that the numbers and specific identity of many rhythmic (phospho)proteins would vary between independently isolated lines due to stochastic heterogeneity and clonal expansion effects, as also occurs with wild type cells ^123,124^. It is also true that the numbers, identities and amplitudes of rhythmic proteins will inevitably vary somewhat, dependent on the method of analysis used to determine rhythmicity (Figure S1B). It is not plausible, however, that a different method of rhythmicity analysis would yield a qualitatively different result. Moreover, our findings have a clear precedent in that oscillations of protein synthesis and degradation stimulate facilitatory metabolism and compensatory ion transport to maintain osmotic and protein homeostasis during the metabolic cycle of yeast cells ^67^. Indeed, the many mechanistic features shared between mammalian circadian rhythms and yeast respiratory oscillations may indicate a common ancestral origin ^125^.

In our kinase inference analysis, we were surprised not to find evidence supportive of circadian phosphorylation by casein kinase 1 or any of the other kinases implicated in the post-translational circadian regulation. Indeed, the very poor overlap in rhythmically phosphorylated proteins between the two genotypes, and between rhythmic phosphorylation and rhythmic protein in either genotype, suggests that the circadian functions of this post-translational modification are likely to be context-dependent. As with the proteome, however, we did notice a highly significant association between change in rhythmicity and change in phosphopeptide abundance. If a similar interpretation to that which we propose for the proteome were true, it would imply that as much as 20% of protein phosphorylation is subject to cell-autonomous circadian regulation (Figure S2D), and in most cases it is matched by a dephosphorylation rhythm of similar phase and amplitude. Given the consolidation of phosphorylation rhythms around a temporal window that anticipates the active phase (Figure 2G), we speculate that the co-ordinated phase-coherent circadian regulation of phosphorylation and dephosphorylation at many phosphosites, without change in their steady state phosphorylation level, might confer more sensitive and rapid transduction of a given extracellular stimulus when received around the rest-to-active transition compared with 12 hours later. Future work will be required to test this hypothesis experimentally.

## Conclusion

In conclusion, we have shown that CRY-dependent feedback mechanisms are not required for cell-autonomous circadian rhythms of protein abundance or phosphorylation in mammalian cells. Moreover, when CRY is present, it functions to suppress more abundance rhythms than it facilitates both *ex vivo* and *in vivo*. CRY-deficiency was associated with an overall imbalance of the proteome, through changes in protein stoichiometries, reduced proteasome activity, increased protein synthesis and overall protein levels. Cells adapt to this genetic insult by altering osmotic balance and proteome composition to achieve a different set point for protein homeostasis, with increased basal stress and sensitivity to proteotoxic stress. This likely accounts for the impairment of circadian timekeeping in CKO cells, tissues and mice, and may also contribute to the (patho)physiological consequences of CRY-deficiency in mice. We speculate that the principal utility of CRY-mediated feedback repression is to couple global protein synthesis with degradation rates for most proteins, to minimise changes in protein abundance. This would ensure the energetically efficient temporal consolidation of proteome renewal during each day, whilst defending protein and osmotic homeostasis.

## Methods

### Mammalian cell culture

All animal work was licensed by the Home Office under the Animals (Scientific Procedures) Act 1986, with Local Ethical Review by the Medical Research Council and the University of Cambridge, UK. Fibroblasts homozygous for PER2::LUCIFERASE ^126^ were extracted from adult mouse lung tissue and then serial passage was used as described previously to induce spontaneous immortalisation ^125,127^. Fibroblasts were cultured in Dulbecco’s Modified Eagle Medium (DMEM), supplemented with 100 units/ml penicillin, 100 µg/ml streptomycin (Gibco) and 10% FetalClone III serum (HyClone, Thermo Fisher). All cells were confirmed to be free of Mycoplasma. Unless stated otherwise, confluent cell cultures up to a maximum of 30 passages were used during experiments to abolish any effects of cell division, since these cells display contact inhibition.

### General statistics

P values are annotated in figures with asterisks, where the number of asterisks indicates the significance: Ns = not significant; * = p≤0.05; ** = p≤0.01, *** = p≤0.001; **** = p≤0.0001. Technical replicates are denoted as “n” in the figures or figure legends (e.g. n=3), and biological replicates are denoted as “N”. Statistical tests were carried out using Prism Graphpad 8 (San Diego, Ca) or R v4.0.3.

### Longitudinal bioluminescent reporter experiments

Data from longitudinal bioluminescence recordings were analysed using Prism Graphpad 8 (San Diego, Ca). A 24-hour moving average was used to detrend data, and a circadian damped cosine wave was fitted by least-squares regression to determine period, phase and amplitude:

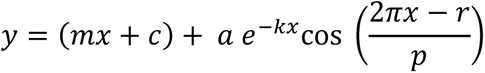

Where *m* is the baseline gradient, *c* is the displacement in the *y* axis, *k* is the damping rate, *a* is the amplitude, *r* is the phase and *p* is the period. The first 24 hours of each recording were omitted because this represents the transient effects of medium change on clock gene expression. Rhythmicity of bioluminescence recordings was assessed by comparing the fit of this equation to the null hypothesis of a straight line using the Extra sum-of-squares F test in Prism Graphpad 8 (San Diego, CA). If fitting to the damped cosine was preferred (p ≤ 0.05) then the recording was deemed “rhythmic”.

### Timecourse experiments: general structure

Cells were plated at a near-confluent density (roughly 27,000 cells per cm^2^) and cultured in DMEM with 10% FetalClone III serum for one week in a temperature-controlled incubator that was programmed to oscillate between 32°C and 37°C, with transitions every 12 hours. The cells received a medium change at the transition between 37°C and 32°C after 4 days. After another 3 days the cells received another medium change at the same transition time into medium containing either 10% or 1% serum, and the incubator was programmed to remain at 37°C constantly. At this time, a subset of cells received medium containing 1 mM luciferin, and these were placed into an ALLIGATOR for bioluminescent recording. After 24 hours, sampling began, with 3 hour intervals, and continuing for 3 days. The time point of the first sample is known as “Experimental time 0”, and all time points are reported relative to this. The nature of the sampling varied according to the specific experiment, and details are presented in separate sections.

### Proteomics and phosphoproteomics

#### Sample preparation

A timecourse was carried out as described above. At each timepoint cells were washed twice in ice cold PBS and then lysed at room temperature in 100 µL lysis buffer (8 M urea, 20 mM Tris, pH 8) for 20 minutes. The lysis buffer was prepared the day before sampling began, and frozen in 1 mL aliquots. At each timepoint, one aliquot was defrosted at room temperature (23°C) whilst shaking at 700 rpm for 5 minutes. After lysis the cells were scraped and technical replicates were combined before flash freezing in liquid nitrogen and storage at −80°C. After defrosting, the samples were sonicated for 2 minutes and the protein concentration was measured using a BCA assay (Pierce). 12 pooled samples were created by combining a portion of each experimental sample such that each sample/pool contained an equal amount of protein. All samples were then flash frozen in liquid nitrogen and stored at −80°C.

#### Enzymatic Digestion

Each sample (256 µg) was reduced with 5 mM DTT at 56°C for 30 minutes and then alkylated with 10 mM iodoacetamide in the dark at room temperature for 30 minutes. They were then digested with mass spectrometry grade Lys-C (Promega) at a protein:Lys-C ratio of 100:1 (w/w) for 4 hours at 25°C. Next, the samples were diluted to 1.5 M urea using 20 mM HEPES (pH 8.5) and digested at 30°C overnight with trypsin (Promega) at a ratio of 70:1 (w/w). Digestion was quenched by the addition of trifluoroacetic acid (TFA) to a final concentration of 1%. Any precipitates were removed by centrifugation at 13000g for 15 minutes. The supernatants were desalted using homemade C18 stage tips containing 3M Empore extraction disks (Sigma) and 5 mg of Poros R3 resin (Applied Biosystems). Bound peptides were eluted with 30-80% acetonitrile (MeCN) in 0.1% TFA and lyophilized.

#### TMT (Tandem mass tag) peptide labelling

The lyophilized peptides from each sample were resuspended in 100 µl of 2.5% MeCN, 250 mM triethylammonium bicarbonate. According to manufacturer’s instructions, 0.8 mg of each TMT 10plex reagent (Thermo) was reconstituted in 41 µl of anhydrous MeCN. The peptides from each time point and pooled sample were labelled with a distinct TMT tag for 75 minutes at room temperature. The labelling reaction was quenched by incubation with 8 µl 5% hydroxylamine for 30 min. For each set of 10-plex TMT reagent, the labelled peptides from 8 time point samples + 2 pools were combined into a single sample and partially dried to remove MeCN in a SpeedVac (Thermo Scientific). After this, the sample was desalted as before and the eluted peptides were lyophilized.

#### Basic pH Reverse-Phase HPLC fractionation

The TMT labelled peptides were subjected to off-line High Performance Liquid Chromatography (HPLC) fractionation, using an XBridge BEH130 C18, 3.5 μm, 4.6 mm x 250 mm column with an XBridge BEH C18 3.5 μm Van Guard cartridge (Waters), connected to an Ultimate 3000 Nano/Capillary LC System (Dionex). Peptide mixtures were resolubilized in solvent A (5% MeCN, 95% 10 mM ammonium bicarbonate, pH 8) and separated with a gradient of 1-90% solvent B (90% MeCN, 10% 10 mM ammonium bicarbonate, pH 8) over 60 minutes at a flow rate of 500 μl/min. A total of 60 fractions were collected. They were combined into 20 fractions and lyophilized and desalted as before. 5% of the total eluate from each fraction was taken out for proteome LC-MS/MS analysis and the rest was used for phosphopeptide enrichment.

#### Enrichment of phosphopeptides

All 20 fractions of peptide mixture were enriched first using PHOS-Select iron affinity gel, an Iron (III) Immobilised Metal Chelate Affinity Chromatography (IMAC) resin (Sigma). Desalted peptides were resuspended in 30% MeCN, 0.25 M acetic acid (loading solution) and 30 µl of IMAC beads, previously equilibrated with the loading solution, was added. After 60 minutes incubation at room temperature, beads were transferred to a homemade C8 (3M Empore) stage tip and washed 3 times with loading solution. Phosphopeptides were eluted sequentially with 0.4 M NH_3_, 30% MeCN, 0.4 M NH_3_ and 20 µl of 50% MeCN, 0.1% TFA.

The flow-through from the C8 stage tips was collected and combined into 10 fractions and used for titanium dioxide (TiO_2_) phosphopeptide enrichment. For this, the total volume of flow-through was made up to 50% MeCN, 2 M lactic acid (loading buffer) and incubated with 1-2 mg TiO_2_ beads (Titansphere, GL Sciences, Japan) at room temperature for 1 hour. The beads were transferred into C8 stage tips, washed in the tip twice with the loading buffer and once with 50% MeCN, 0.1% TFA. Phosphopeptides were then eluted sequentially with 50 mM K_2_HPO_4_ (pH 10) followed by 50% MeCN, 50 mM K_2_HPO_4_ (pH 10) and 50% MeCN, 0.1% TFA.

The first 10 fractions of IMAC and the 10 fractions of TiO_2_ enriched phosphopeptides were combined, and the other 10 fractions from IMAC enrichment were combined into 5 fractions, thus making a total of 15 fractions for phosphoproteomics analysis. Phosphopeptide solution from these fractions were acidified, partially dried, and desalted with a C18 Stage tip that contained 1.5 µl of Poros R3 resin. These were then partially dried again and thus ready for mass spectrometry analysis.

#### LC MS/MS

The fractionated peptides were analysed by LC-MS/MS using a fully automated Ultimate 3000 RSLC nano System (Thermo) fitted with a 100 μm x 2 cm PepMap100 C18 nano trap column and a 75 μm × 25 cm reverse phase C18 nano column (Aclaim PepMap, Thermo). Samples were separated using a binary gradient consisting of buffer A (2% MeCN, 0.1% formic acid) and buffer B (80% MeCN, 0.1% formic acid), and eluted at 300 nL/min with an acetonitrile gradient. The outlet of the nano column was directly interfaced via a nanospray ion source to a Q Exactive Plus mass spectrometer (Thermo). The mass spectrometer was operated in standard data-dependent mode, performing a MS full-scan in the m/z range of 350-1600, with a resolution of 70000. This was followed by MS2 acquisitions of the 15 most intense ions with a resolution of 35000 and Normalised Collision Energy (NCE) of 33%. MS target values of 3e6 and MS2 target values of 1e5 were used. The isolation window of precursor ion was set at 0.7 Da and sequenced peptides were excluded for 40 seconds.

#### Spectral processing and peptide and protein identification

The acquired raw files from LC-MS/MS were processed using MaxQuant (Cox and Mann) with the integrated Andromeda search engine (v1.6.3.3). MS/MS spectra were quantified with reporter ion MS2 from TMT 10plex experiments and searched against the *Mus musculus* UniProt Fasta database (Dec 2016). Carbamidomethylation of cysteines was set as fixed modification, while methionine oxidation, N-terminal acetylation and phosphorylation (STY) (for phosphoproteomics group only) were set as variable modifications. Protein quantification requirements were set at 1 unique and razor peptide. In the identification tab, second peptides and match between runs were not selected. Other parameters in MaxQuant were set to default values.

The MaxQuant output file was then processed with Perseus (v1.6.2.3). Reporter ion intensities were uploaded to Perseus. The data was filtered: identifications from the reverse database were removed, only identified by site, potential contaminants were removed, and we only considered proteins with ≥1 unique and razor peptide. Then all columns with an intensity “less or equal to zero” were converted to “NAN” and exported. The MaxQuant output file with phosphor (STY) sites table was also processed with Perseus software (v1.6.2.3). The data was filtered: identifications from the reverse database were removed, only identified by site, potential contaminants were removed and we only considered phosphopeptides with localization probability ≥ 0.75. Then all columns with intensity “less or equal to zero” were converted to “NAN” and exported.

#### Bioinformatics

All data handling was done using R v3.6.3. Since the sample for timepoint 12 was missing for CRY1^-/-^; CRY2^-/-^, abundance values were inferred for each protein by taking the mean of the two neighbouring timepoints. WT and CKO datasets were analysed either combined or independently. The combined analysis was used to directly compare protein and phosphopeptide abundances between genotypes, since the internal reference scaling normalisation accounts for batch effects. The independent method was used for all other analysis that did not require comparison of abundance, thus allowing the detection of proteins that were present in one genotype but not the other.

Proteins and phosphopeptides were only accepted for further analysis if present in all timepoints and pooled samples. Hence in the combined analysis, proteins/phosphopeptides had to be present in all timepoints for both genotypes, as well as all pooled samples. In the independent analysis, proteins/phosphopeptides had to be present in all timepoints and pools for one genotype only. Sample loading normalisation was carried out by taking the sum of all intensities for each time point and normalising to the mean of these, since an equal amount of protein was used for each TMT labelling reaction. This was followed by internal reference scaling (IRS) to allow for comparisons between TMT experiments ^128^: for each TMT 10plex set the mean abundance for each protein in both pools was calculated. Then the mean of these means was calculated and used to normalise the values for each protein for all the samples.

Rhythmicity was tested using the RAIN (Rhythmicity Analysis Incorporating Non-parametric methods) algorithm ^129^, and multiple testing was corrected for using the adaptive Benjamini-Hochberg method. Proteins with a corrected p ≤ 0.05 were deemed significant. Relative amplitude of rhythmic proteins was calculated by detrending the data using a 24-hour moving average and dividing the resultant range by the average normalised protein abundance. To include only proteins with a biologically relevant level of oscillation, only those with relative amplitude ≥ 10% were taken for further analysis (see text for details). Phosphoproteomics data were handled in the same way, except that normalised phosphopeptide abundances were adjusted according to the changes in abundance of the corresponding protein from the normalised total proteome data, and no threshold for relative amplitude was used.

Gene ontology analysis was performed using the GOrilla online tool ^130,131^. Analysis was performed either as a single ranked list of gene names, or as a target dataset compared to background (all proteins detected in the experiment). Kinase recognition motifs were screened using a custom script written in Python v2.7, which used the PHOSIDA database ^52,53^. Scripts and processed data are available on GitHub (https://github.com/davwong47/Circadian-proteomics). Mass spectrometry data have been deposited to the ProteomeXchange Consortium ^132^ via the PRIDE ^133^ partner repository with the dataset identifier PXD019499.

### Western blotting

For Western blots, proteins were run on NuPAGE^TM^ Novex^TM^ 4-12% Bis-Tris Protein Gels (Thermo Fisher) before transferring to nitrocellulose membranes. For transfer, the iBlot system (Thermo Fisher) was used. Membranes were blocked using 5% milk powder (Marvel) or 0.125% BSA (Sigma) and 0.125% milk powder (Marvel) in TBS containing 0.1% Tween-20 (TBST) for 30 minutes at room temperature then incubated with primary antibody at 4°C overnight. HRP-conjugated secondary antibodies (Thermo Fisher) diluted 1:10000 in blocking buffer were incubated with the blots for 1 hour at room temperature. Chemiluminescence was detected in a Biorad chemidoc using Immobilon reagent (Millipore). Protein loading was checked by staining gels with Colloidal Coomassie Blue Stain (Severn Biotech). Densitometric analysis was carried out using Image Lab 4.1 (Biorad Laboratories 2012).

### Measurement of cellular protein content

At specified time points, confluent monolayers of cells were washed twice with ice-cold PBS. Cells were then incubated with 200 µL digitonin lysis buffer (50 mM Tris pH 7.4, 0.01% digitonin, 5 mM EDTA, 150 mM NaCl, 1 U/mL Benzonase, protease and phosphatase inhibitors) on ice for 15 minutes before lysates were collected. For total protein extraction, cells were instead incubated with 200 µL RIPA buffer (50 mM Tris pH 7.4, 1% SDS, 5 mM EDTA, 150 mM NaCl, 1 U/mL Benzonase, protease and phosphatase inhibitors), on ice for 15 minutes. Cells lysed with RIPA buffer were then scraped and collected and all samples were flash frozen in liquid nitrogen. After thawing, RIPA lysates were sonicated at high power for 10 seconds at 4°C to shear genomic DNA. RIPA lysates and digitonin lysates were clarified by centrifugation at 21,000 g for 15 minutes at 4°C.

Intrinsic tryptophan fluorescence was used to measure protein concentrations. 10 µL of each sample was transferred into a UV-transparent 384 well plate (Corning 4681) in quadruplicate. After brief centrifugation of the plate, quantification was carried out using a Tecan Spark 10M microplate reader, with excitation at 280 nm and emission at 350 nm. Standards were made using bovine serum albumin (Fisher Scientific), dissolved using the same lysis buffer as the lysates being measured. Standard curves were fitted to a quadratic curve using Prism Graphpad 8 (San Diego, Ca), and protein concentrations were interpolated.

### Measurement of intracellular ion content by Inductively Coupled Plasma - Mass Spectrometry (ICP-MS)

Confluent monolayers of cells were washed on ice with iso-osmotic buffer A (300 mM sucrose, 10 mM Tris base, 1 mM EDTA, pH 7.4 adjusted with phosphoric acid, 330-340 mOsm adjusted with sucrose/HPLC water), followed by iso-osmotic buffer B (300 mM sucrose, 10 mM Tris base, 1 mM EDTA, pH 7.4 adjusted with acetic acid, 330-340 mOsm adjusted with sucrose/HPLC water). Iso-osmotic buffer A contains phosphoric acid which displaces lipid bound ions. Iso-osmotic buffer B contains acetic acid which removes traces of phosphates. Cells were then incubated for 30 minutes at room temperature in 200 µL ICP-MS cell lysis buffer (65% nitric acid, 0.1 mg/mL (100 ppb) cerium). Lysates were then collected and stored at −80°C. All samples were thawed simultaneously and diluted using HPLC water to a final concentration of 5% nitric acid. Diluted samples were analysed by Inductively Coupled Plasma - Mass Spectrometry (ICP-MS) using the NexION 350D ICP-MS (PerkinElmer Inc.) as described previously ^134^.

### Proteasome activity assays

20,000 cells per well were plated in a 96-well plate using standard culture medium, and on the following day the medium was changed. 10 µM epoxomicin in the medium was used as negative control. 3 hours later, the ProteasomeGlo Cell-Based Assay (Promega) was used to measure proteasome catalytic activity. Chymotrypsin-like, trypsin-like and caspase-like activities were measured separately using the relevant substrates from Promega (Suc-LLVY-Glo, Z-LRR-Glo, Z-nLPnLD-Glo respectively). Assay reagents were prepared according to the manufacturer’s instructions. The 96-well plate was equilibrated to room temperature, and a volume of assay reagent equal to the volume of medium was added to each well before shaking at 700 rpm for 2 minutes. The plate was incubated at room temperature for a further 10 minutes, and then luminescence was measured using the Tecan Spark 10M microplate reader, recording counts for 1 second. The luminescence readings from the epoxomicin controls represent background protease activity, and so this was subtracted from all other recordings.

### Measurement of translation rate

WT and CKO mouse lung fibroblasts were grown to confluence in 48-well plates, and medium was changed 24h before the experiment to either low (1%) or high (10%) serum. The cells were pulsed with 0.1 mCi/ml ^35^S-L-methionine/^35^S-L-cysteine mix (EasyTag™ EXPRESS35S Protein Labeling Mix, Perkin Elmer) in cysteine/methionine-free DMEM for 15 min at 37°C, with or without serum supplement. Afterwards, cells were washed with ice-cold PBS and lysed in digitonin-based buffer (with protease inhibitor tablet, added freshly) on ice. Lysates were reduced with LDS buffer and run on 4-12% Bis-Tris SDS-PAGE using MES buffer. Gels were then dried at 80°C and exposed overnight to a phosphorimager screen. Images were acquired with Typhoon FLA700 gel scanner and quantified using Fiji.

For the puromycin pulse experiment, cells were seeded (in the presence of 1 mM luciferin) in fibronectin-coated dishes at high density one day before starting the experiment. Separate 6-well plates were used for each time point. 30 minutes before the start of the experiment, cells were synchronised by a dexamethasone pulse (10 nM), after which cells were exchanged into Air medium (DMEM without glucose, L-glutamine, phenol red, sodium pyruvate and sodium bicarbonate (Sigma). The following were added: 0.35 g/L sodium bicarbonate (Sigma), 5 g/L glucose (Sigma), 20 mM MOPS (VWR), penicillin/streptomycin solution (as above), Glutamax (Thermo Fisher), B27 (Thermo Fisher), 1 mM potassium luciferin solution (Biosyth), 1% FetalClone III serum. The medium was adjusted to pH 7.6 at room temperature, and osmolality adjusted to 350-360 mOsm/kg using sodium chloride). At the indicated time points, 10 µg/mL puromycin was added while keeping cells warm on a hot plate, after which the plate was incubated in tissue culture incubator for exactly 10 minutes. Timepoint 0 represents cells just before the dexamethasone pulse. Cells were then washed with ice-cold PBS, after which they were lysed in lysis buffer (50 mM tris, pH 7.5, 1% Triton X-100, 0.1% SDS, 1.5 mM MgCl_2_, 100 mM NaCl, cOmplete protease inhibitor cocktail (Roche)) and flash frozen. For Western blot analysis, samples were spun down, diluted in Laemmli sample buffer, ran on 4-20% gradient gels, and blots were probed for ATF4 and tubulin.

### Animal work

All animals were maintained in 12:12 hour light:dark conditions. Post-mortem histopathology examinations were carried out by Abbey Veterinary Services, UK. For the *in vivo* stress marker experiment, animals received an IP injection of bortezomib (Generon) (2.5 mg/kg) or vehicle control (1% DMSO in sterile PBS) 2 hours after the transition to the light phase. 5 hours later, the animals were culled and lungs were immediately flash frozen in liquid nitrogen. Tissues were homogenised on ice in RIPA buffer (as above) and flash frozen. After thawing these samples on ice, they were sonicated briefly at 4°C. Samples were clarified by centrifugation and protein diluted to 2 mg/mL. Bolt/NuPage loading buffer with 50 mM TCEP (final concentration) was added and reduced for 10 minutes at 70°C. Western blots were then carried out as described above, probing against phospho-STAT3 (Tyr705) or actin. One female mouse and one male mouse of each genotype were used in this experiment.

### Antibodies

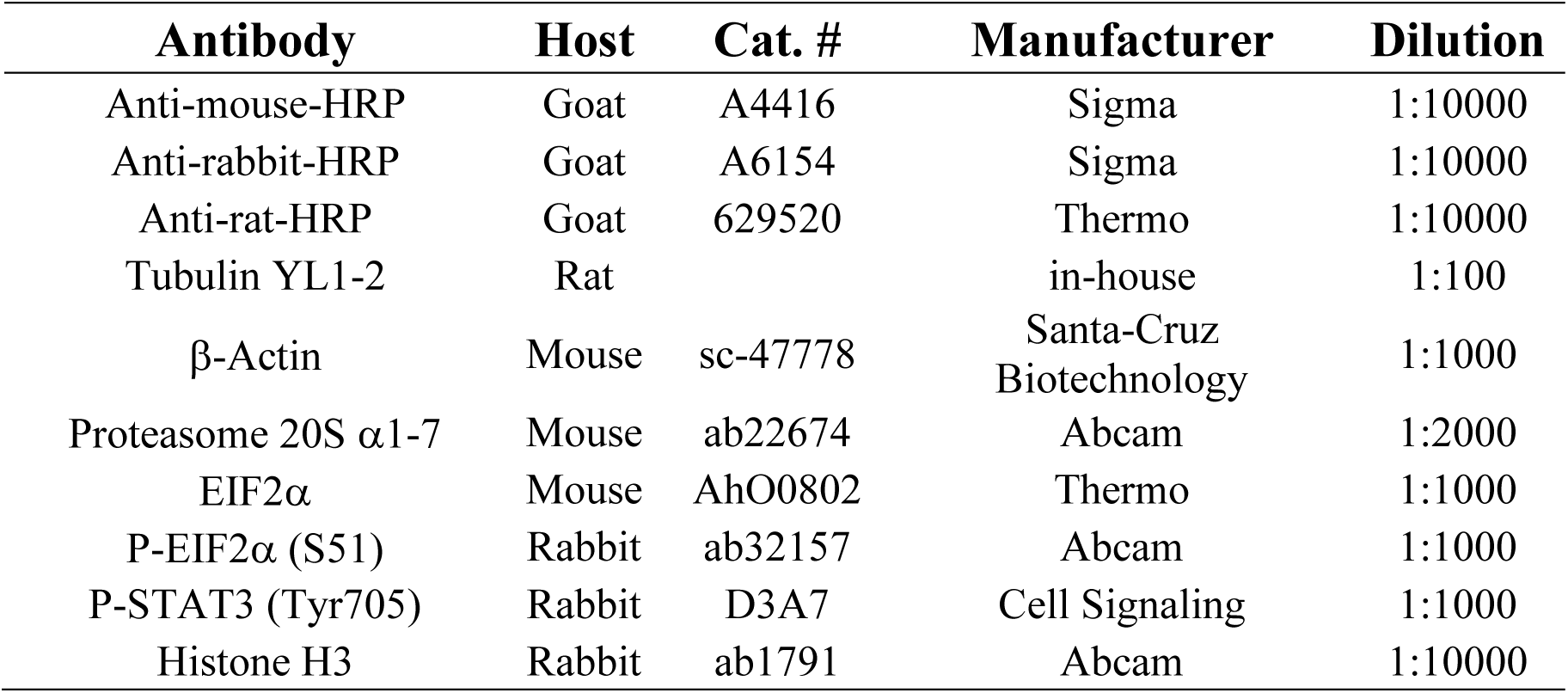

## Acknowledgements

We thank biomedical technical staff at Medical Research Council (MRC) Ares facility and LMB facilities for assistance; G.T. van der Horst and J.S. Takahashi for sharing rodent models; Y-G. Suh, M.H. Hastings and E.S. Maywood for providing reagents and input; M. Hegde, E. Zavodszky, R. Edgar, N. Hoyle, M. Coetzee, J. Chesham, A. Hufnagel, P. Crosby, D.S. Tourigny, J.E. Chambers, H.C. Causton and Tim Stevens for assistance with analysis and experiments as well as valuable discussion. DCSW was supported by the MRC Doctoral Training Programme and the Frank Edward Elmore Fund. AS was supported by the AstraZeneca Blue Skies Initiative. NMR was supported by the Medical Research Council (MR/S022023/1). MP was supported by the Dutch Cancer Foundation (KWF, BUIT-2014-6637) and EMBO (ALTF-654-2014). JON was supported by the Medical Research Council (MC_UP_1201/4) and the Wellcome Trust (093734/Z/10/Z).

## Contributions

DCSW, ES and JON designed the study, analysed the data and wrote the manuscript; DCSW, ES, AS, AZ and MP performed cell experiments; SPC and JD performed mass spectrometry analyses; DCSW, NMR, ADB and JON performed mouse studies; MR performed tissue collection and husbandry. All authors commented on the manuscript.

