## Supplementary information for "CRYPTOCHROME suppresses the circadian proteome and promotes protein homeostasis"

#### Supplementary figure legends

**Supplementary Figure 1, Relating to Figure 1. Examples and analysis of (phospho)proteomics. a)** Post-hoc analysis of proteomics data from Mauvoisin et al. 2014 [1]. Volcano plot showing the fold change in average expression of all proteins, in CKO livers compared to WT (q = Benjamini-Hochberg corrected p-value). Statistically significant changes ($q \leq 0.05$) are shown in red. Some proteins are labelled as space allows. **b)** The relative abundance of CRY1 detected in the WT dataset is plotted. It was preferentially fit by a damped cosine wave over a straight line (Extra sum-of-squares F test, p=0.01). Below: Longitudinal bioluminescence recording of WT PER2::LUC fibroblasts, performed simultaneously with the proteomics experiment as a phase marker. Mean±SEM.

**c)** RAIN and eJTK were compared as statistical tests for rhythmicity. Top, the results of eJTK analysis showed that the relative numbers of rhythmic proteins and the overlap were similar to the results from the RAIN analysis in Figure 1. **d), e)** Heatmaps showing min-max normalised plots for all the rhythmic proteins (d) and phosphopeptides (e) in WT and CKO cells. For each genotype separately, rows represent proteins sorted by phase, and each column is a time point from the timecourse experiment. One missing sample is annotated in black. **f), g)** Examples of proteins (f) and phosphopeptides (g) detected by mass spectrometry are shown, labelled by gene name and phosphosite. Examples are included of proteins/phosphopeptides that are rhythmic in both genotypes, rhythmic in only one genotype, or not rhythmic in either. P-values shown are from a comparison of fit (F test, damped cosine against straight line). Annotation as rhythmic/arrhythmic are from the RAIN output.

**Supplementary Figure 2, Relating to Figures 1 & 2. Summaries of (phospho)proteomics analysis. a)** Table summarising the numbers of proteins detected in both genotypes with significantly increased or decreased abundance (corrected $p<0.05$ vs. $p\geq0.05$) and significant change in rhythmicity (RAIN $p<0.05$ vs. $p\geq0.05$) between WT and CKO cells. If it is assumed that

CRY is required for canonical TTFL function, then rhythms in the abundance of more proteins are suppressed by the TTFL (green) than are dependent upon it (blue), and the abundance of most detected proteins changes as a consequence of CRY deletion. There was a significant association between change in abundance and change in rhythmicity (Fisher's exact test, $p=0.04$). **b)** Using all proteins detected in our proteomics experiment using *ex vivo* constant conditions, fold change was calculated from maximum/minimum protein abundances. Individual protein abundances are plotted. Mann-Whitney test: $p<0.0001$. **c)** Data was extracted from Mauvoisin et al. [1], where the proteome was quantified in WT and CKO mouse livers extracted under diurnal conditions. Fold change was calculated from maximum/minimum protein abundances. Individual protein abundances are plotted. Mann-Whitney test: $p<0.0001$. **d)** Table summarising the numbers of phosphopeptides detected in both genotypes with significantly increased or decreased abundance (corrected $p<0.05$ vs. $p\geq0.05$) and significant change in rhythmicity (RAIN $p<0.05$ vs. $p\geq0.05$) between WT and CKO cells. If it is assumed that CRY is required for canonical TTFL function then more rhythms in protein phosphorylation are suppressed by the TTFL (green) than are facilitated by it (blue), and most detected protein phosphorylation changes as a consequence of CRY deletion. There was a significant association between change in abundance and change in rhythmicity (Fisher's exact test, $p=0.009$). **e)** Using all phosphopeptides detected in our proteomics experiment using *ex vivo* constant conditions, fold change was calculated from maximum/minimum protein abundances. Individual protein abundances are plotted. Mann-Whitney test: $p<0.0001$.

**Supplementary figure S3, Relating to Figures 1 & 2. Protein phosphorylation is increased and protein phosphatase expression reduced in CKO compared with WT cells. a)** Fold-change in abundance was calculated for each phosphopeptide. There was a significant upregulation of phosphorylation in CKO cells compared to WT (One sample t test, $p<0.0001$, $n=2803$). **b)** Fold-change in abundance was calculated for protein kinases (top) and protein phosphatases (bottom) detected in the proteomics dataset. There was a significant downregulation of phosphatase abundance in CKO cells compared to WT (One sample t test, p=0.002, n=39), but no overall change in kinase abundance (One sample t test, p=0.7, n=182). **c), d), e)** Using the phosphoproteomics dataset, phosphopeptide sequences were analysed, with the number of kinase binding motifs counted for a panel of 25 kinases present in the PHOSIDA database [2,3]. Phosphopeptides that were rhythmic (in WT, in CKO, or in both genotypes respectively) were compared to the background of phosphopeptides present in all samples and pools.

**Supplementary Figure 4, Relating to Figure 3. Increased protein synthesis in CKO cells. a)** Whole images of Western blots shown in Figure 3B, probing for alpha subunits of the 20S proteasome (left) and Histone H3 (right). Samples were taken at 12h and 36h after a medium change. Only samples taken at 36h are shown in Figure 3B because those from 12h are likely to represent an acute response to the medium change. **b)** $^{35}$S-methionine incorporation was used to measure translation rate in cultured WT and CKO cells. This was carried out at 0% and 10% serum. 4 replicates are shown, run on the same gel. An image of the phosphor screen is shown above, with the corresponding Coomassie stain below. The condition with 10% serum is shown in Figure 3E as this represents normal culture conditions. **c)** Quantification of (A). 2-way ANOVA showed interaction between the effects of serum and genotype (p=0.045). Holm-Sidak multiple comparisons results are shown as asterisks. N=3 experiments, n=4 technical replicates.

**Supplementary Figure 5, Relating to Figure 4. Increased rhythmic ion transporter expression and activity in CKO compared with WT cells. a), b)** 29 proteins were annotated as "Ion transport" by the GO analysis from the proteomics experiment – relative amplitude and average abundance was calculated for each of these proteins, in both WT and CKO. On average, both relative amplitude and average abundance was increased in CKO cells compared to WT (Paired t test). **c)** Examples of ions detected from both WT and CKO cells by ICP-MS are shown.

Mean±SEM, F test for comparison of fit between damped cosine and straight line, N=3. The preferred fit is shown in red.

**Supplementary Figure 6, Relating to Figure 5. Increased stress in CKO cells and reduced growth/viability in CKO mice. a)** Volcano plot showing the fold change in average expression of all proteins in CKO cells compared to WT (*q* = Benjamini-Hochberg corrected p-value). Proteins annotated as "Response to stress" from GO analysis are highlighted in red, showing that these are upregulated in CKO cells. **b), c)** Quantification from proteomics results of average DENR and eIF2A abundance. Mean±SD, t test with Welch correction. **d), e)** Growth curves of male and female mice were weighed weekly. Mice were of the following genotypes: WT, CRY1$^{-/-}$; CRY2$^{+/-}$ and CRY1$^{-/-}$; CRY2$^{-/-}$ (CKO). F test was used to test the null hypothesis that one curve fits all sets. P values annotated as asterisks. **f)** Food consumption measured over 1 week, normalised for mouse weight. Food consumption was monitored by weighing food daily. Mean±SD, 2-way ANOVA. **g)** Death rates among the 3 genotypes mentioned above, expressed as a percentage of the number of mice. Mice were fed *ad libitum* and group-housed under standard 12:12 light:dark conditions. The absolute numbers of deaths and total population size are annotated on the bars. Only mice that had been weaned were included, and unnatural causes of death (e.g. cage flooding, fighting) were excluded. Asterisk indicates significance from Chi-squared test for trend, p=0.007. Comparing WT and CKO, Fisher's exact test p=0.009. Comparing WT and Het, Fisher's exact test p=0.5. Median age of death for WT mice = 4 weeks, CRY1$^{-/-}$; CRY2$^{+/-}$ = 4 weeks, CRY1$^{-/-}$; CRY2$^{-/-}$ (CKO) = 7 weeks, p=0.4 for differences between groups, Kruskal-Wallis test.

**Figure S7, Relating to Figure 6. A refined model for the generation and utility of cellular circadian timekeeping that accounts CKO phenotypes. a)** The canonical model for circadian timekeeping and the effect of the CRY1$^{-/-}$;CRY2$^{-/-}$ (CKO) genotype. The transcriptional-translational feedback loop (TTFL) is the mechanistic basis by which circadian rhythms in clock protein activity are generated. This leads to rhythmic expression of clock-controlled genes and encoded proteins, ultimately facilitating the circadian coordination of cell biology and whole-organism physiology; depicted by sinusoidal curves in black. CRY proteins are essential components of the TTFL, and without them it cannot function. In consequence, there is no circadian regulation of cellular function, organismal physiology or behaviour (flat line, red). Absence of circadian rhythms leads to the deleterious phenotypes of CKO mice and cells, such as altered metabolism, increased carcinogenic potential and death.

**b)** A refinement to the canonical model where blue elements are present in both WT and CKO, black elements in WT only, and red elements are consequences of $CRY1^{-/-}$;$CRY2^{-/-}$ (CKO) genotype. We suggest the principal utility of the canonical TTFL is to minimise circadian rhythms in protein abundance, not to generate them, in order to allow daily cycles of protein activity and proteome renewal whilst maintaining protein homeostasis overall. In this model, circadian timing results from a cytosolic post-translational oscillator, or "cytoscillator", involving enzymes such as casein kinase I as in mammalian erythrocytes and other eukaryotic cells. The cytoscillator is sufficient to confer post-translational circadian regulation upon protein activity, abundance, and compensatory ion transport, but its timing mechanism is not robust against proteotoxic and other external stresses. PERIOD proteins are the primary vector of timing information from the cytosol to the nucleus. The cytoscillator confers daily rhythms upon the activity of PERIOD as well as CRY, BMAL1 and other promiscuous transcription/translation factors proteins, which act to anticipate and buffer against changes in protein abundance, both directly and *via* clock-controlled gene regulation. This gives rise to circadian regulation of chromatin architecture and the expression of genes including *Period1/2* and *Cry1/2* seen in WT cells; and confers robustness to circadian rhythms by preventing proteome imbalance and proteotoxic stress, as well as by hysteresis *via* PERIOD1/2 abundance rhythms [4]. In consequence, many proteins are circadian regulated in their activity and synthesis, but relatively few proteins show biologically significant change in their overall abundance. Cells adapt to the absence of CRY by remodelling of the (phospho)proteome and ionome, at the cost of increased cellular stress, which underlies the many adverse phenotypes of the CKO genetic model. One such adverse phenotype is a severe impairment to the temporal co-ordination of physiology and behaviour over circadian timescales. CKO cells possess the capacity for circadian timekeeping but, without the TTFL to temporally buffer protein homeostasis, epigenetic adaptations to CRY-deletion are effectively epistatic to the expression of circadian rhythms *in vivo* under most conditions. This model is an extension of that proposed by Putker et al (2021)[5] and is consistent with the vast majority of data of which we are aware. In future work, amongst other predictions from this model, we will explore the evidence for circadian regulation of compartmentalisation and proteome renewal.

FIGURE S1

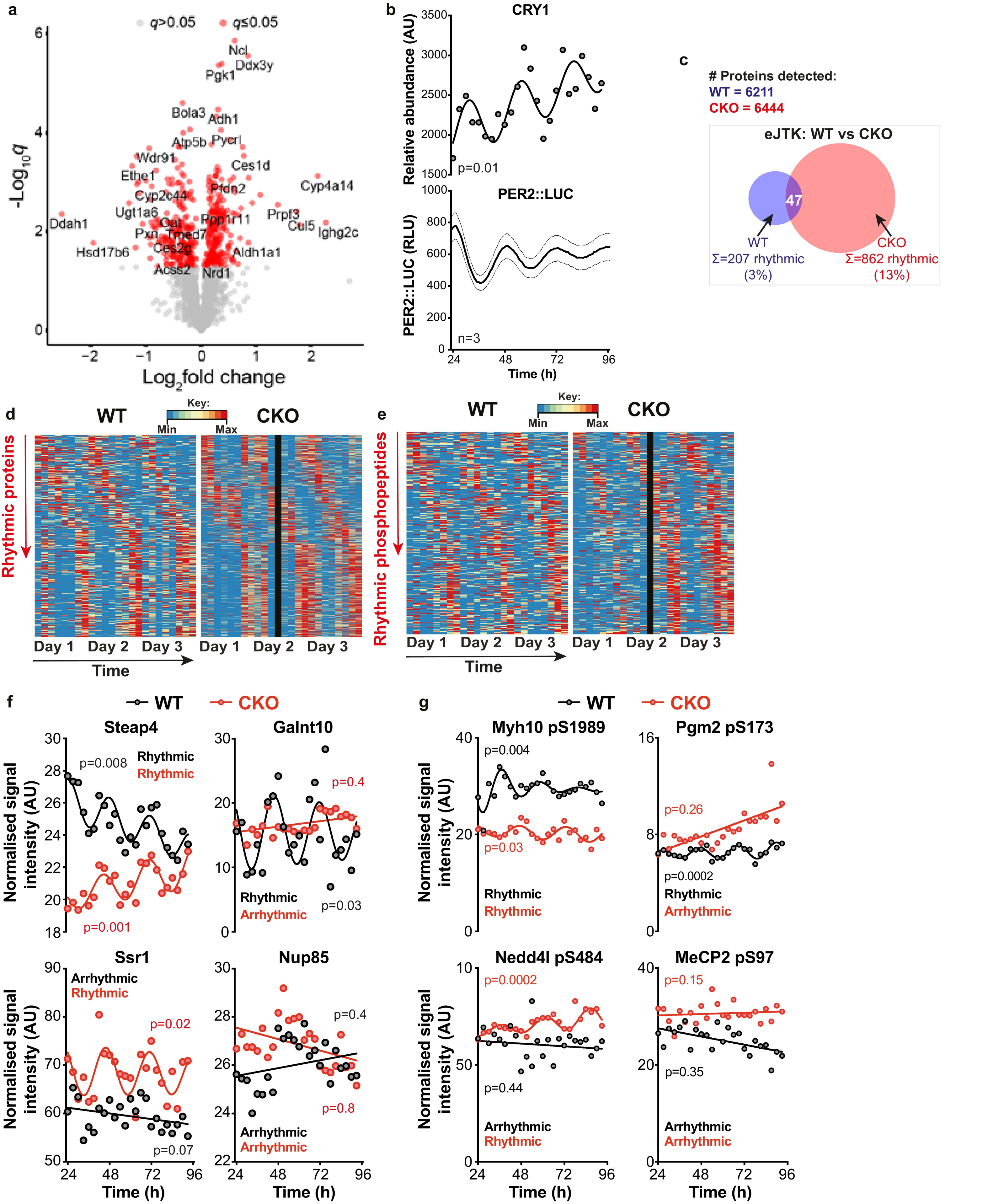
a
q>0.05
q≤0.05
-Log10q
Log2fold change
Ncl
Ddx3y
Pgk1
Bola3
Adh1
Atp5b
Pycrl
Wdr91
Ces1d
Ethe1
Cyp4a14
Cyp2c44
Pfdn2
Ugt1a6
Prpf3
Ddah1
Ppp1r11
Cul5
Ighg2c
Oat
Pxn
Tmed7
Hsd17b6
Ces2g
Aldh1a1
Acss2
Nrd1
b
CRY1
Relative abundance (AU)
p=0.01
PER2::LUC
PER2::LUC (RLU)
n=3
Time (h)
c
### Proteins detected:
WT = 6211
CKO = 6444
eJTK: WT vs CKO
47
WT
Σ=207 rhythmic
(3%)
CKO
Σ=862 rhythmic
(13%)
d
WT
Key:
Min
Max
CKO
Rhythmic proteins
Day 1
Day 2
Day 3
Day 1
Day 2
Day 3
Time
e
WT
Key:
Min
Max
CKO
Rhythmic phosphopeptides
Day 1
Day 2
Day 3
Day 1
Day 2
Day 3
Time
f
WT
CKO
Steap4
p=0.008
Rhythmic
Rhythmic
p=0.001
Galnt10
p=0.4
Rhythmic
Arrhythmic
p=0.03
Normalised signal intensity (AU)
Ssr1
Arrhythmic
Rhythmic
p=0.02
p=0.07
Nup85
p=0.4
p=0.8
Arrhythmic
Arrhythmic
Time (h)
g
WT
CKO
Myh10 pS1989
p=0.004
p=0.03
Rhythmic
Rhythmic
Pgm2 pS173
p=0.26
p=0.0002
Rhythmic
Arrhythmic
Nedd4l pS484
p=0.0002
p=0.44
Arrhythmic
Rhythmic
MeCP2 pS97
p=0.15
p=0.35
Arrhythmic
Arrhythmic
Normalised signal intensity (AU)
Time (h)

FIGURE S2

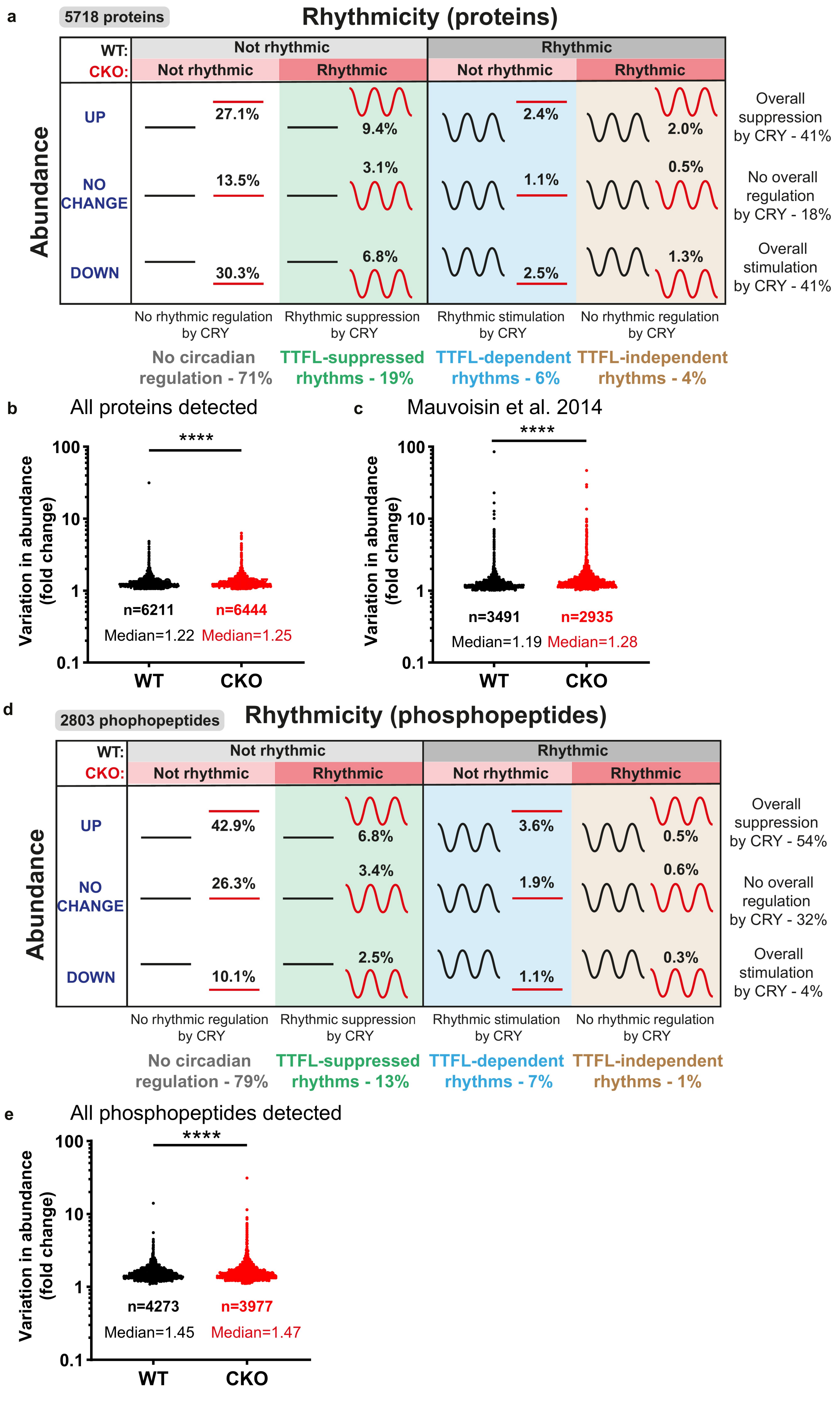

a
5718 proteins
Rhythmicity (proteins)
WT:
CKO:
Not rhythmic
Rhythmic
Not rhythmic
Rhythmic
Not rhythmic
Rhythmic
Abundance
UP
NO CHANGE
DOWN
27.1%
13.5%
30.3%
9.4%
3.1%
6.8%
2.4%
1.1%
2.5%
2.0%
0.5%
1.3%
Overall suppression by CRY - 41%
No overall regulation by CRY - 18%
Overall stimulation by CRY - 41%
No rhythmic regulation by CRY
Rhythmic suppression by CRY
Rhythmic stimulation by CRY
No rhythmic regulation by CRY
No circadian regulation - 71%
TTFL-suppressed rhythms - 19%
TTFL-dependent rhythms - 6%
TTFL-independent rhythms - 4%
b
All proteins detected
****
Variation in abundance (fold change)
100
10
1
0.1
n=6211
n=6444
Median=1.22
Median=1.25
WT
CKO
c
Mauvoisin et al. 2014
****
Variation in abundance (fold change)
100
10
1
0.1
n=3491
n=2935
Median=1.19
Median=1.28
WT
CKO
d
2803 phophopeptides
Rhythmicity (phosphopeptides)
WT:
CKO:
Not rhythmic
Rhythmic
Not rhythmic
Rhythmic
Not rhythmic
Rhythmic
Abundance
UP
NO CHANGE
DOWN
42.9%
26.3%
10.1%
6.8%
3.4%
2.5%
3.6%
1.9%
1.1%
0.5%
0.6%
0.3%
Overall suppression by CRY - 54%
No overall regulation by CRY - 32%
Overall stimulation by CRY - 4%
No rhythmic regulation by CRY
Rhythmic suppression by CRY
Rhythmic stimulation by CRY
No rhythmic regulation by CRY
No circadian regulation - 79%
TTFL-suppressed rhythms - 13%
TTFL-dependent rhythms - 7%
TTFL-independent rhythms - 1%
e
All phosphopeptides detected
****
Variation in abundance (fold change)
100
10
1
0.1
n=4273
n=3977
Median=1.45
Median=1.47
WT
CKO

FIGURE S3

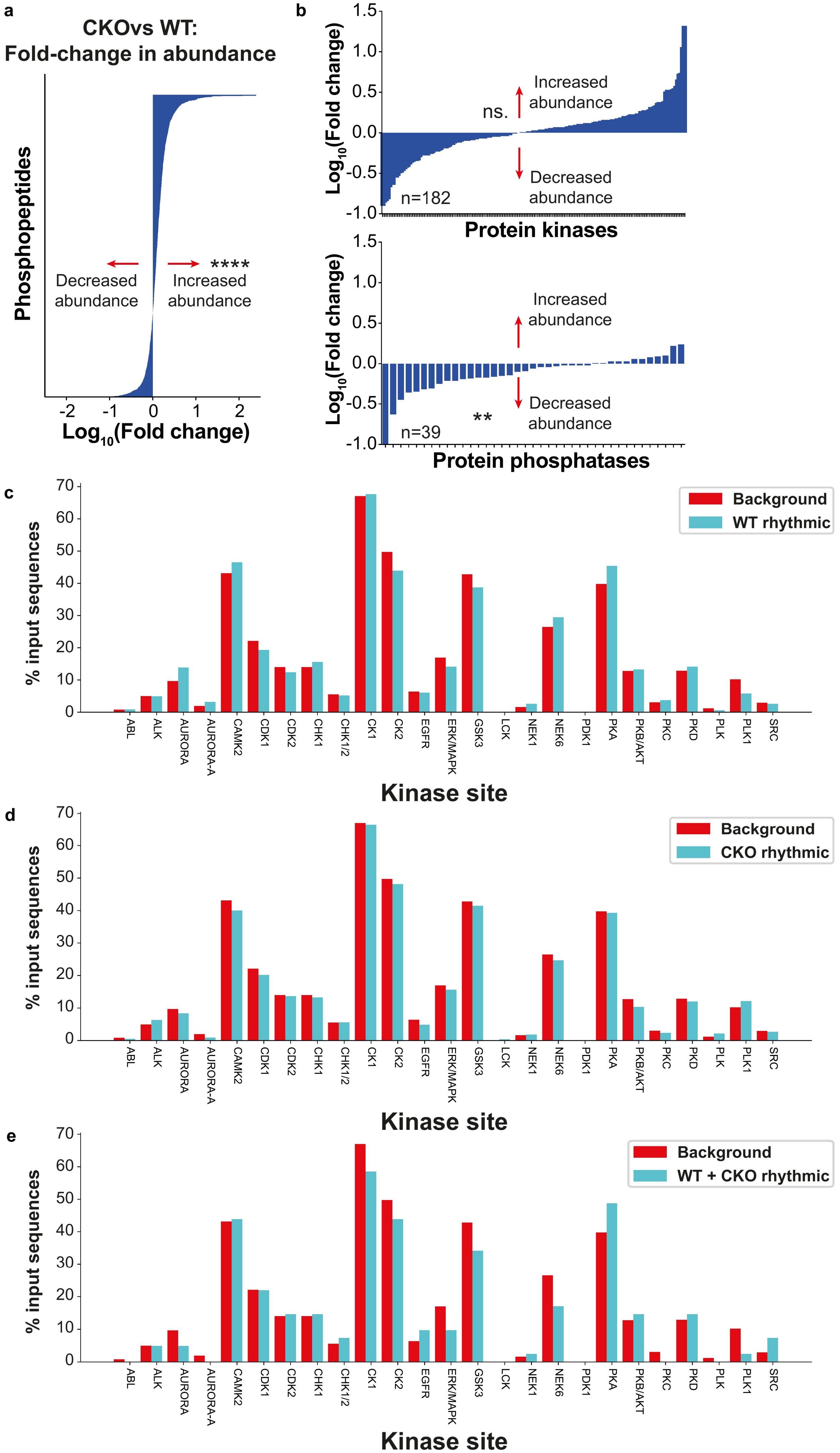
a
CKOvs WT:
Fold-change in abundance
Phosphopeptides
Decreased abundance
Increased abundance
****
Log10(Fold change)
b
Log10(Fold change)
ns.
Increased abundance
Decreased abundance
n=182
Protein kinases
Increased abundance
Decreased abundance
**
n=39
Protein phosphatases
c
% input sequences
Background
WT rhythmic
ABL
ALK
AURORA
AURORA-A
CAMK2
CDK1
CDK2
CHK1
CHK1/2
CK1
CK2
EGFR
ERK/MAPK
GSK3
LCK
NEK1
NEK6
PDK1
PKA
PKB/AKT
PKC
PKD
PLK
PLK1
SRC
Kinase site
d
Background
CKO rhythmic
Kinase site
e
Background
WT + CKO rhythmic
Kinase site

FIGURE S4

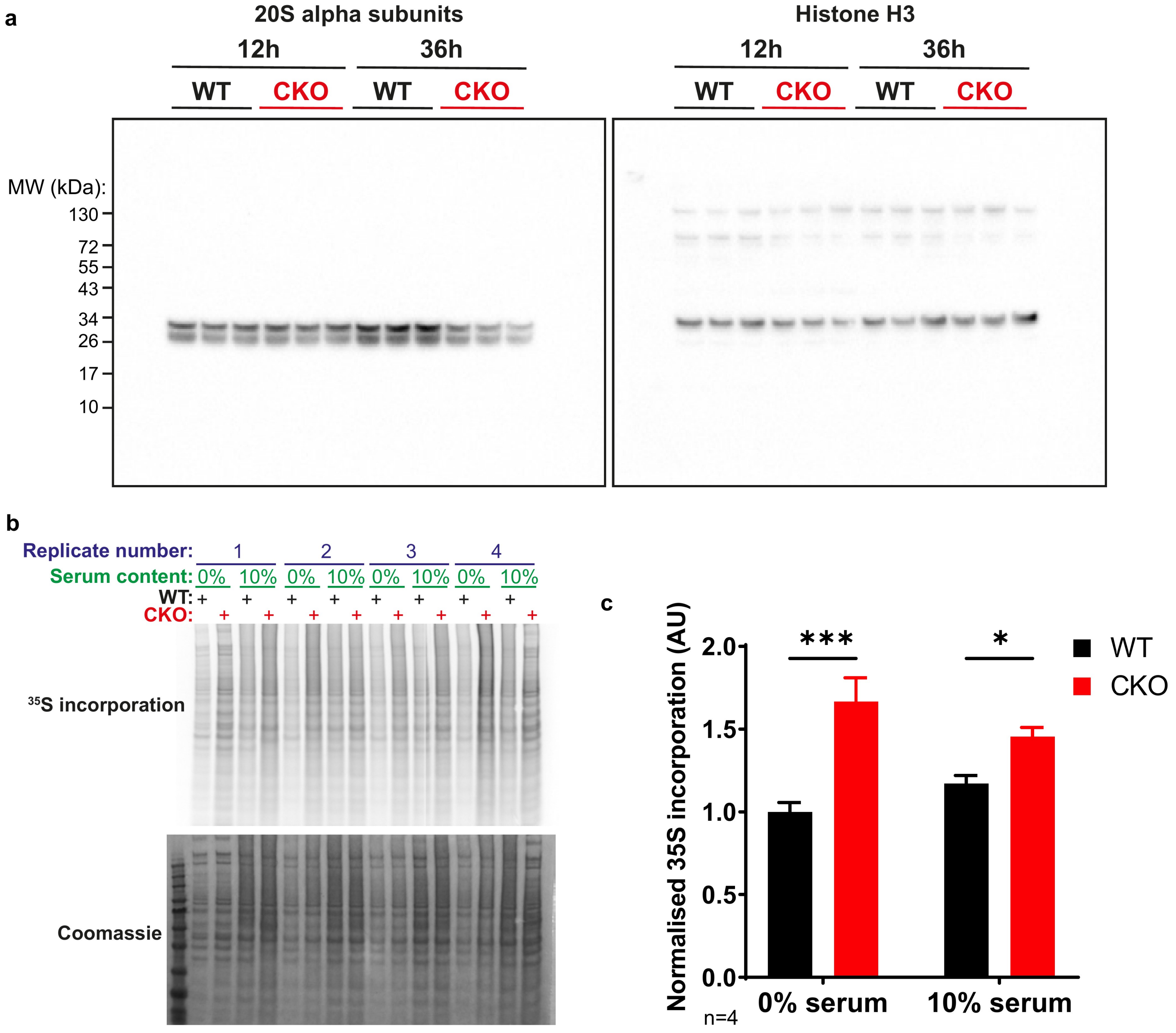

a
20S alpha subunits
Histone H3
12h
36h
WT
CKO
MW (kDa):
130
72
55
43
34
26
17
10
b
Replicate number:
1
2
3
4
Serum content:
0%
10%
WT:
CKO:
35S incorporation
Coomassie
c
Normalised 35S incorporation (AU)
2.0
1.5
1.0
0.5
0.0
***
*
WT
CKO
0% serum
10% serum
n=4

### FIGURE S5

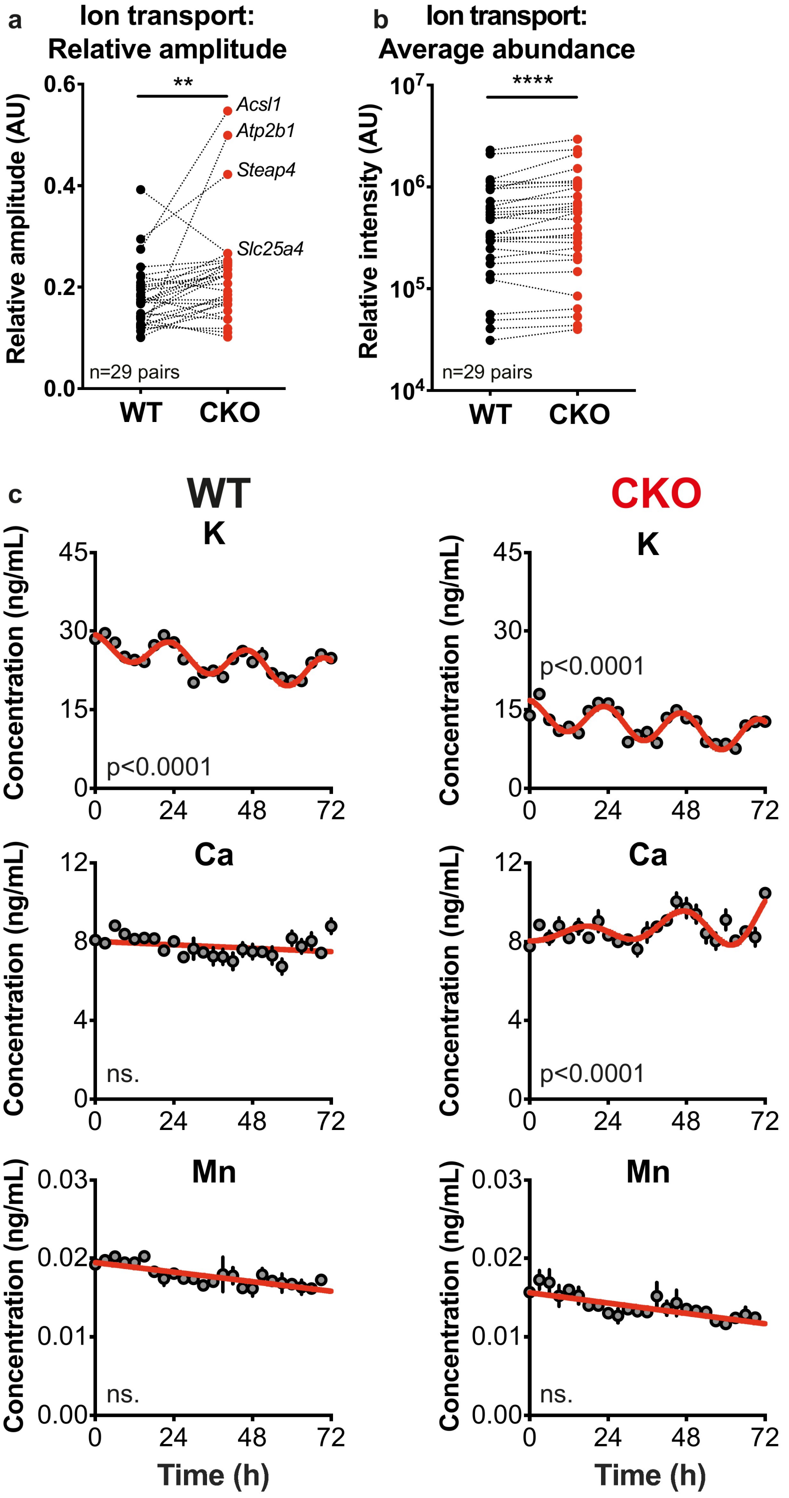
a
Ion transport:
Relative amplitude
Relative amplitude (AU)
0.6
0.4
0.2
0.0
**
Acsl1
Atp2b1
Steap4
Slc25a4
n=29 pairs
WT
CKO
b
Ion transport:
Average abundance
Relative intensity (AU)
10^7
10^6
10^5
10^4
****
n=29 pairs
WT
CKO
c
WT
CKO
K
Concentration (ng/mL)
45
30
15
0
p<0.0001
0
24
48
72
Ca
12
8
4
0
ns.
p<0.0001
Mn
0.03
0.02
0.01
0.00
ns.
Time (h)

FIGURE S6

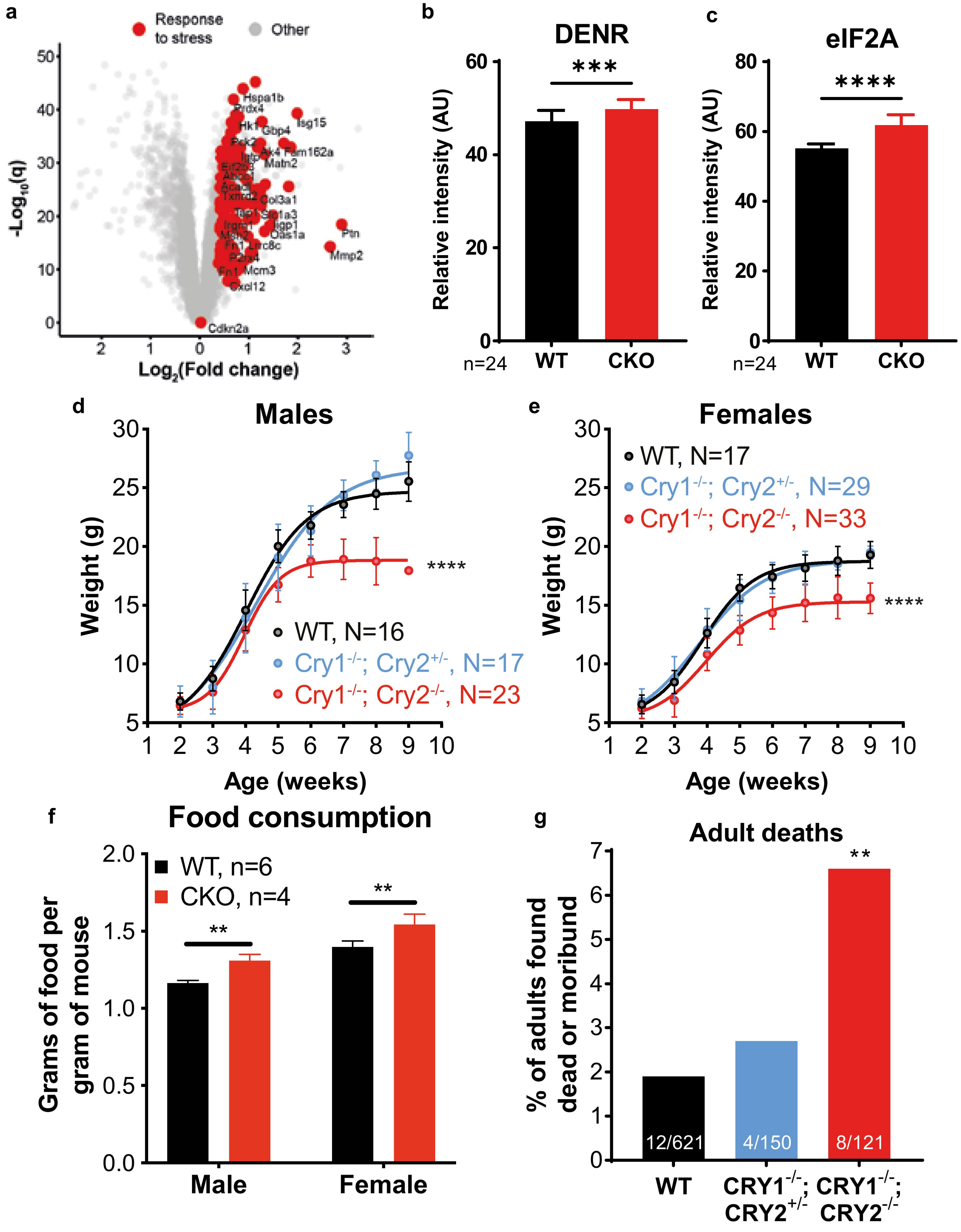
a
Response to stress
Other
-Log10(q)
Log2(Fold change)
Hspa1b
Prdx4
Isg15
Hk1
Gbp4
Pck2
Ak4
Fam162a
Matn2
Col3a1
Slc1a3
Iigp1
Oas1a
Ptn
Mmp2
Fn1
Lrrc8c
P2rx4
Mcm3
Cxcl12
Cdkn2a
b
DENR
***
Relative intensity (AU)
n=24
WT
CKO
c
eIF2A
****
Relative intensity (AU)
n=24
WT
CKO
d
Males
Weight (g)
Age (weeks)
****
WT, N=16
Cry1-/-; Cry2+/-, N=17
Cry1-/-; Cry2-/-, N=23
e
Females
WT, N=17
Cry1-/-; Cry2+/-, N=29
Cry1-/-; Cry2-/-, N=33
Weight (g)
Age (weeks)
****
f
Food consumption
WT, n=6
CKO, n=4
Grams of food per gram of mouse
**
**
Male
Female
g
Adult deaths
% of adults found dead or moribund
**
12/621
4/150
8/121
WT
CRY1-/-; CRY2+/-
CRY1-/-; CRY2-/-

FIGURE S7

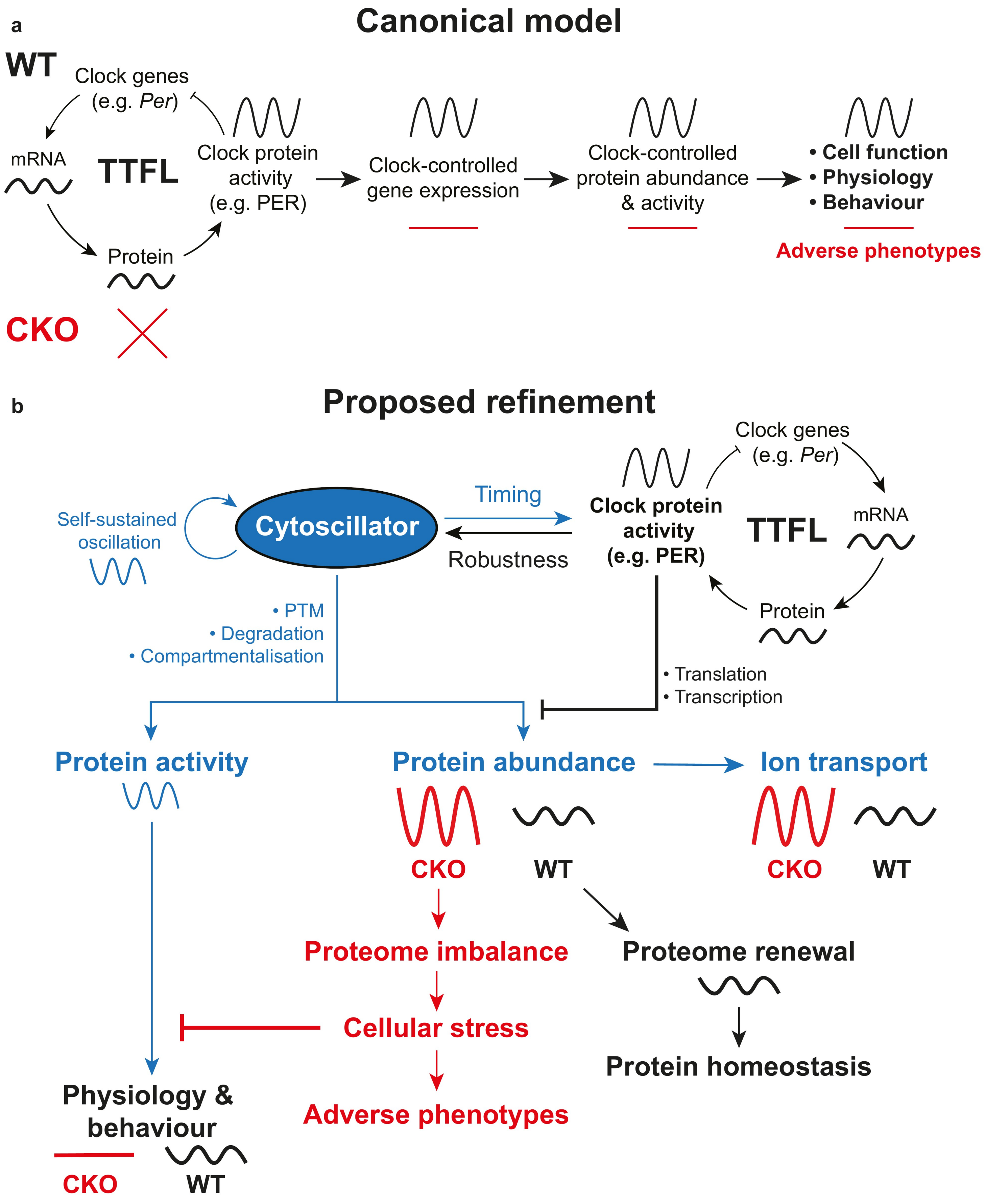

a
Canonical model
WT
Clock genes
(e.g. *Per*)
mRNA
TTFL
Protein
Clock protein
activity
(e.g. PER)
Clock-controlled
gene expression
Clock-controlled
protein abundance
& activity
• Cell function
• Physiology
• Behaviour
Adverse phenotypes
CKO
b
Proposed refinement
Self-sustained
oscillation
Cytoscillator
Timing
Robustness
Clock genes
(e.g. *Per*)
Clock protein
activity
(e.g. PER)
TTFL
mRNA
Protein
• PTM
• Degradation
• Compartmentalisation
• Translation
• Transcription
Protein activity
Protein abundance
Ion transport
CKO
WT
CKO
WT
Proteome imbalance
Proteome renewal
Cellular stress
Protein homeostasis
Physiology &
behaviour
Adverse phenotypes
CKO
WT
